# RNA encodes physical information

**DOI:** 10.1101/2024.12.11.627970

**Authors:** Ian Seim, Vita Zhang, Ameya P. Jalihal, Benjamin M. Stormo, Sierra J. Cole, Joanne Ekena, Hung T. Nguyen, D. Thirumalai, Amy S. Gladfelter

## Abstract

Most amino acids are encoded by multiple codons, making the genetic code degenerate. Synonymous mutations affect protein translation and folding, but their impact on RNA itself is often neglected. We developed a genetic algorithm that introduces synonymous mutations to control the diversity of structures sampled by an mRNA. The behavior of the designed mRNAs reveals a physical code layered in the genetic code. We find that mRNA conformational heterogeneity directs physical properties and functional outputs of RNA-protein complexes and biomolecular condensates. The role of structure and disorder of proteins in biomolecular condensates is well appreciated, but we find that RNA conformational heterogeneity is equally important. This feature of RNA enables both evolution and engineers to build cellular structures with specific material and responsive properties.

---

The degeneracy of the genetic code allows for multiple codons to be exchanged in an mRNA coding sequence and still produce the same polypeptide (*1, 2*). This degeneracy can result in a combinatorial explosion such that 10^86^ possible mRNA sequences can encode a 200 amino acid protein. Analysis of codon usage variation has focused on protein production and folding rates (*3, 4*), yet codons also directly influence RNA structures, making synonymous mutations far from silent in terms of the RNA polymer itself. RNA structure controls RNA-binding protein recruitment (*5*), subcellular localization (*6*), RNA editing (*7*), and stability (*8*). In many of these contexts, RNAs are components of biomolecular condensates (*9*).

Many condensates are enriched in RNA-binding proteins with low complexity sequences (LCS) that are predicted to contain intrinsically disordered regions (IDRs) (*10*). IDR conformational heterogeneity supports multivalent interactions, and specific residues influence the composition and physical properties of condensates (*11, 12*). In contrast to proteins, how conformational heterogeneity or disorder in RNA polymers influences the properties of condensates has not been systematically examined. We set out to measure how mutations in RNA sequences that are synonymous for the purpose of protein coding alter the conformational heterogeneity of RNAs and impact the physical properties of mesoscale cellular assemblies. We show that the primary sequence of mRNAs encodes physical information and that this embedded code can exploit synonymous mutations to impact RNA function.

## Algorithmic design of RNA structural ensembles using *in silico* evolution with synonymous mutations

Due to the small number of nucleotides and its hydrophilic nature, RNA is inherently conformationally promiscuous and can sample many energetically comparable structures, commonly resulting in more rugged and shallow energy landscapes compared to globular proteins (*13*). Therefore, conformational heterogeneity is far more descriptive of RNA structure than a single, minimum free energy state (*14*). The predicted conformational heterogeneity of sequences can be approximated by the ensemble diversity (ED), which is the Boltzmann-weighted average pairwise distance between all secondary structures in the ensemble (*15*). Distance is defined as the number of base pairs needed to rearrange to transform one secondary structure into another. The ED can vary greatly between RNA sequences, and different mRNAs with distinct EDs have been correlated with altered morphology of biomolecular condensates (*16*). However, the physical mechanisms by which RNA ED impacts mesoscale cell assemblies like condensates remain unexplored, in striking contrast to protein counterparts. A challenge in examining how ED impacts RNA assemblies lies in isolating the property of ED from the many other features and functions of RNA molecules.

We found an opportunity to address this problem when we noticed that ED can vary substantially even for mRNAs encoding the same protein when comparing sequences between individuals of the same species (**Fig 1A**). Specifically, there is a wide range of ED within the cyclin mRNA *CLN3* from wild isolates of the fungus *Ashbya gossypii* (*17*) (**Fig 1A**). Interestingly, these ED differences arise in large part from a significant enrichment of synonymous mutations (**Fig S1**) and prompted us to predict that synonymous mutations may be important for generating different EDs for a given mRNA, even within a species (*18*). *CLN3* forms biomolecular condensates through interactions with an RNA-binding protein called Whi3. *CLN3*-Whi3 condensates are a well-established model for examining RNA-driven condensate formation *in vitro* and in cells (*19, 20*). We set out to design *CLN3* mRNA sequences with widely variable ED that arises solely from synonymous mutations to examine how RNA conformational heterogeneity impacts condensates.

**Figure 1:**
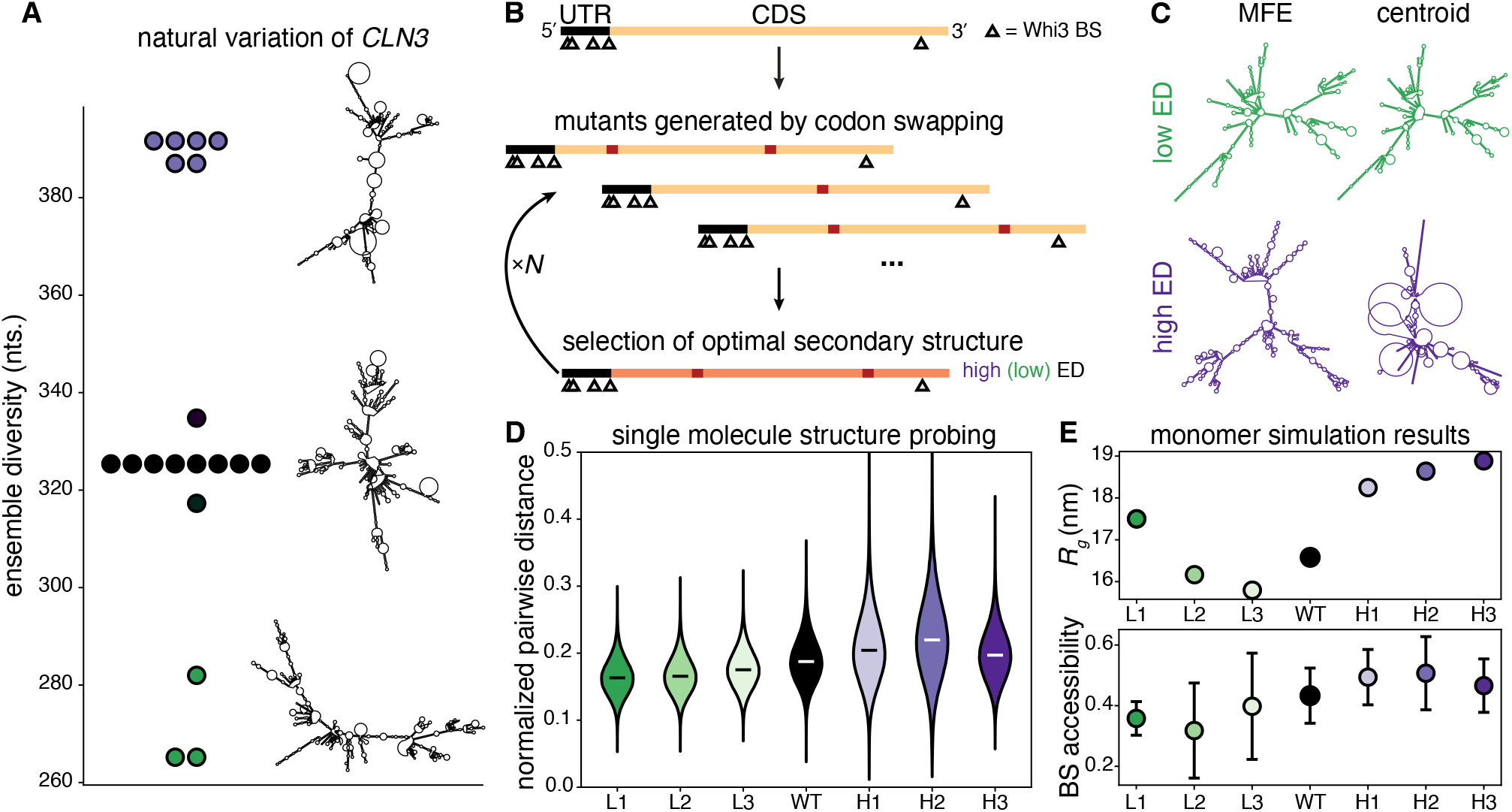
Design of RNA structural ensembles. **(A)** *CLN3* mRNAs in *Ashbya gossypii* wild isolates have different sequences and sample a range of predicted ensemble diversities (ED). Representative centroid structures are shown to the right of the plot points. **(B)** Top: A schematic of the *CLN3* transcript is shown in which the lengths of the 5′ UTR and the coding sequence (CDS) and the positions of the Whi3 binding sites (BS) are drawn to scale. Middle: Several mutant sequences are generated by random synonymous mutations within the CDS by swapping codons subject to the constraints described in the text. Bottom: The mutant sequence with the maximum (minimum) predicted ED is chosen as the parent of the next generation. This process is iterated *N* times until a sequence with desired properties is found. **(C)** Secondary structure predictions of the minimum free energy (MFE) and centroid structures from designed sequences L3 (top) and H3 (bottom) at 25°C and 150mM NaCl are shown. **(D)** The normalized pairwise distances among the longest 1000 reads for each sequence are shown. White and black bars represent medians. All distributions are significantly different based on the Mann-Whitney U test with p < 0.01. **(E)** Top: Predicted radius of gyration (*R*_*g*_) from simulations for each sequence. Error bars are smaller than markers. Bottom: Predicted average Whi3 binding site (BS) solvent accessibility from simulations for each sequence. Error bars represent standard deviations of accessibility among the 5 BS for each sequence.

We created a genetic algorithm that designs mutants of a given mRNA sequence using codon swapping within the coding sequence that preserves the encoded protein. mRNA sequences are selected for maximum (or minimum) predicted ensemble diversity (ED) as determined by RNAfold (*15*) (**Fig 1B, S2**). We initialized the algorithm with the sequence of *CLN3* from the lab reference strain. Mutations in each generation are accepted subject to the following constraints: transcript length is unchanged, the UTRs are unchanged, nucleotide composition is within 1% of the reference sequence, usage of each codon is within 10% of codon usage in the reference sequence, encoded amino acid sequence is unchanged, and no Whi3 binding sites (BS) are created or destroyed. We designed 3 different *CLN3* sequences with minimized predicted ED (L1, L2, L3) and 3 sequences with maximized predicted ED (H1, H2, H3). Multiple sequences of each class were designed to generalize ED as the physical property of focus and eliminate sequence-specific phenomena. The sequence identity varies similarly within and between the different L and H designs, eliminating sequence bias concerns (**Fig S3**). Despite the strict design constraints, we are still able to access highly distinct conformational ensembles. For the designed sequence, ED is visually apparent in the differences or similarities between the predicted minimum free energy (MFE) structure, and the centroid structure, which is the structure closest to all others in the ensemble (**Fig 1C**).

We experimentally validated that the evolved sequences behaved as modeled by *in vitro* transcribing WT *CLN3* and the 6 structure mutants and using RNA structure probing with single-molecule, long read, direct RNA sequencing using Nanopore technology (*21*). This approach enabled us to measure the pairwise distance between the structures of individual mRNA molecules in the population. Any adenosines in single-stranded regions can be chemically modified and this adduct state is read out in single-read RNA sequencing. To estimate the ED of each sequence, we calculated all of the pairwise distances among the 1000 longest reads, normalized by their degree of overlap (**Fig S4**). Low ED sequences (L1, L2, L3) have lower pairwise distances, indicating structures that are more similar to each other, and high ED sequences (H1, H2, H3) have higher pairwise distances, indicating structures that less similar to each other (**Fig 1D**). The average Jensen-Shannon distance (JSD) among the pairwise distance distributions for the low ED sequences is 0.11, among the high ED distributions is 0.2, and between the high and low ED distributions is 0.39 (**Fig S5**). Unprobed control sequences show no differences in pairwise distances, but distance measurements of predicted structural ensembles for these sequences agree with the probed data (**Fig S5, S6**). We find a Pearson’s correlation coefficient between predicted length-normalized ED (NED) and average pairwise distances of 0.89 and between average pairwise distances from the unprobed control and NED of −0.03 (**Fig S7**). We also mathematically show an inverse linear relationship between ED and the variance of nucleotide structural states across a given sequence and observe this relationship experimentally and for predicted structures (**Supplementary Text** and **Fig S8**). Importantly, both the ED-pairwise distance and ED-variance relations are independent of the accuracy of the prediction of any given secondary structure but rather reflect inherent features of the structural ensemble. Although the inaccuracy of predictions of specific secondary structures for long RNAs has been shown (*22*), these data experimentally validate that the ensemble-level designed structures indeed have different ED and behave as predicted.

To generate predictions about 3-dimensional structures, we performed Langevin dynamics simulations of each sequence as monomers using a previously published RNA forcefield (*23*). Low ED sequences were predicted to have smaller radii of gyration (*R*_*g*_) and less solvent-accessible Whi3 binding sites (BS) than their high ED counterparts which may promote higher protein recruitment (**Fig 1E**). The radius of gyration and ED predictions for each sequence are consistent with native RNA gels (**Fig S9**), further validating that these RNA sequences indeed behave as designed to have high or low ED.

## RNA structural heterogeneity controls RNA-protein complex size

To characterize the effects of ED on RNA assemblies, we first analyzed single molecules of the different *CLN3* mutants with total internal reflection fluorescence (TIRF) microscopy. In subsaturated conditions (*24*), a single phase is formed by *CLN3* molecules clustering with Whi3 protein (**Fig 2A**). High ED *CLN3* mutants form larger and brighter *CLN3* puncta than low ED *CLN3* mutants, indicating more recruitment of RNA by high ED sequences (**Fig 2A, B**). The average JSD among the low ED *CLN3* intensity distributions is 0.12, among the high ED distributions is 0.2, and between the low and high ED distributions is 0.33 (**Fig S10**). There is little apparent colocalization with protein because a very low percentage of Whi3 protein was labeled and few molecules comprise a given cluster. Protein is, however, required to facilitate significant interactions among *CLN3* molecules as experiments with a similar concentration of RNA with no Whi3 protein in the same buffer conditions reveals smaller clusters that do not vary in size among *CLN3* structural mutants (**Fig S10, S11**).

**Figure 2:**
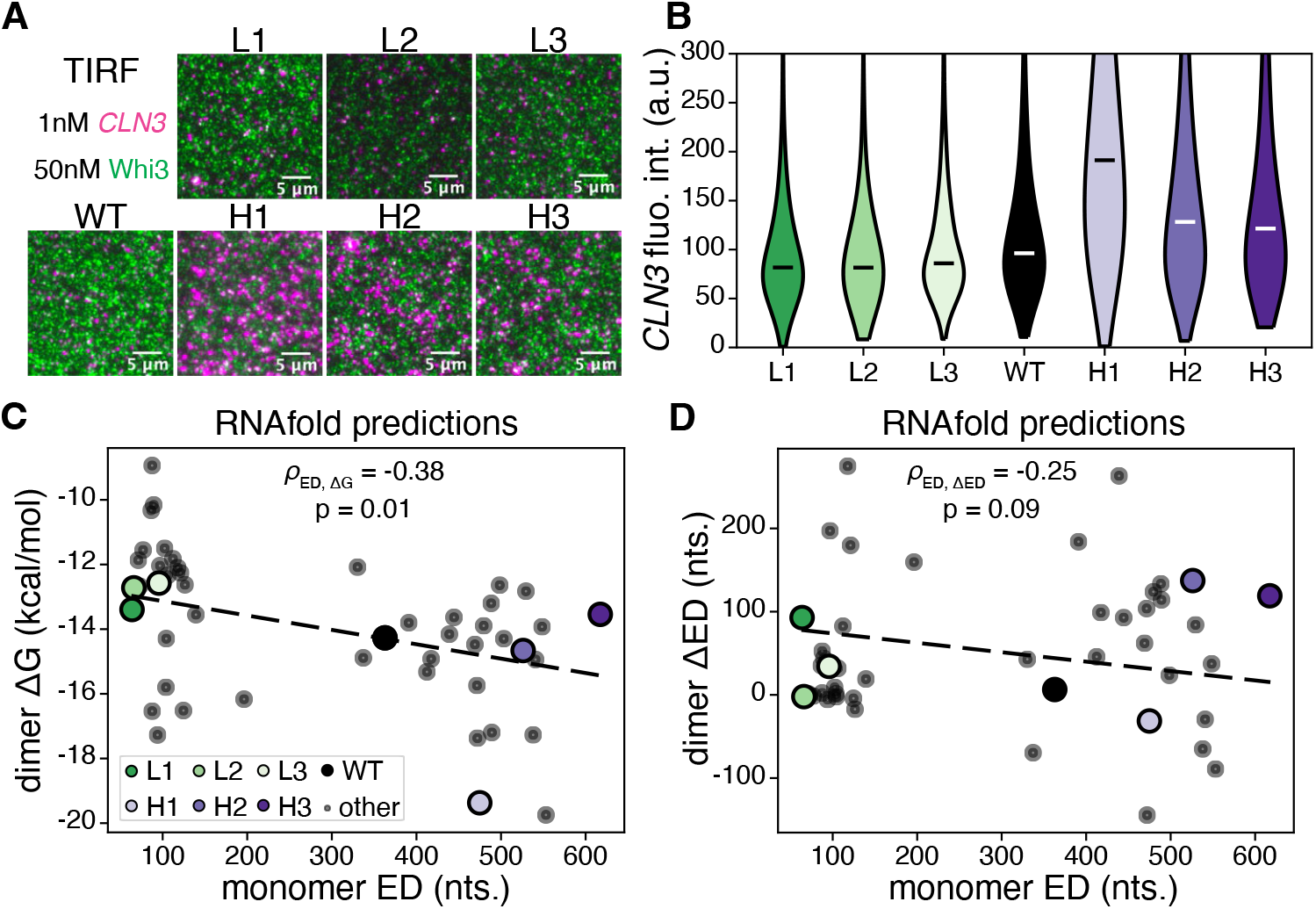
RNA ED controls composition of subsaturated RNA-protein assemblies. **(A)** Total internal reflection fluorescence (TIRF) microscopy was performed to visualize subsaturated clusters of *CLN3* structural mutants and Whi3 protein. Magenta corresponds to *CLN3*, and green corresponds to Whi3. Scale bars are 5μm. **(B)** Distributions of puncta intensities from the data represented in panel (A) are shown. White and black bars indicate medians. All distributions are significantly different based on the Mann-Whitney U test with p < 0.01. **(C)** RNAfold predictions of dimer ΔG, defined in the text, versus monomer ED, for the *CLN3* structure mutants and 40 additional designed sequences (“other” in legend) are shown. *ρ*_ED, ΔG_ is the Pearson’s correlation coefficient, and p is the associated p-value. **(D)** The same sequences are analyzed as in (C) but for dimer ΔED and monomer ED. *ρ*_*ED*,*ΔED*_ is the Pearson’s correlation coefficient between monomer ED and dimer ΔED.

To investigate the connection between ED and cluster formation, we modeled RNA homodimer conformational ensembles for each sequence using RNAcofold, which predicts the free energy of the structural ensemble, *G*, and the ED for dimers (*25*). We computed changes to *G* and ED upon dimerization by comparing their values for the dimer with the values for 2 copies of non-interacting monomers, which gives a change in *G* or ED associated with dimerization, dimer ΔG = G_dimer_ – 2 × G_monomer_, and similarly for dimer ΔED. Additionally, we designed 40 new sequences in order to investigate general relationships between monomer ED and homodimer properties. We found that high ED sequences on average had a more negative predicted *dimer ΔG* compared to low ED sequences, indicating more favorable dimer formation (**Fig 2C**), especially the H1 mutant which also showed the greatest RNA recruitment (**Fig 2B**). The prediction that dimer formation is more energetically favorable for the high ED *CLN3* mutants than the low ED mutants is consistent with the patterns of ED-dependent *CLN3* recruitment to subsaturated clusters. Interestingly, we also observed that dimers formed by high ED sequences on average gained less ED upon dimerization than their low ED counterparts, although the trend was weak (**Fig 2D**). This behavior suggests possible differences in conformational entropy costs upon dimerization, or higher-order associations, between the structure mutants.

## Biomolecular condensate material properties and function controlled by RNA structural heterogeneity

We next examined the impact of RNA conformational heterogeneity on mesoscopic condensate formation. In higher bulk concentrations that promote *CLN3* and Whi3 phase separation, the low ED *CLN3* mutants form large spherical droplets with Whi3, while the high ED *CLN3* mutants form extensive branched networks that appear to be dynamically arrested clusters of very small droplets (**Fig 3A, C**). In addition to different shapes, condensate composition varies, with low ED *CLN3* mutants recruiting both more *CLN3* and Whi3 than their high ED counterparts, and WT recruiting intermediate amounts across a range of combinations of bulk concentrations (**Fig 3B, S12**). The ability of low ED *CLN3* mutants to recruit more protein molecules to the dense phase upon phase separation despite their lower predicted Whi3 BS accessibility (**Fig 1D**) is in striking contrast to the behavior seen at subsaturated conditions (**Fig 2A, B**) and suggests different assembly properties across the phase boundary. The morphological differences likely reflect underlying differences in the viscoelasticity of the condensates (*26*), with low ED *CLN3* mutants forming liquid-like condensates and high ED *CLN3* mutants forming elastic, dynamically arrested condensates. These findings show that RNA conformational heterogeneity can inform the composition and material state of condensates, and that RNAs of the same length, nucleotide composition, and protein binding site valence can generate highly distinct types of assemblies at different scales.

**Figure 3:**
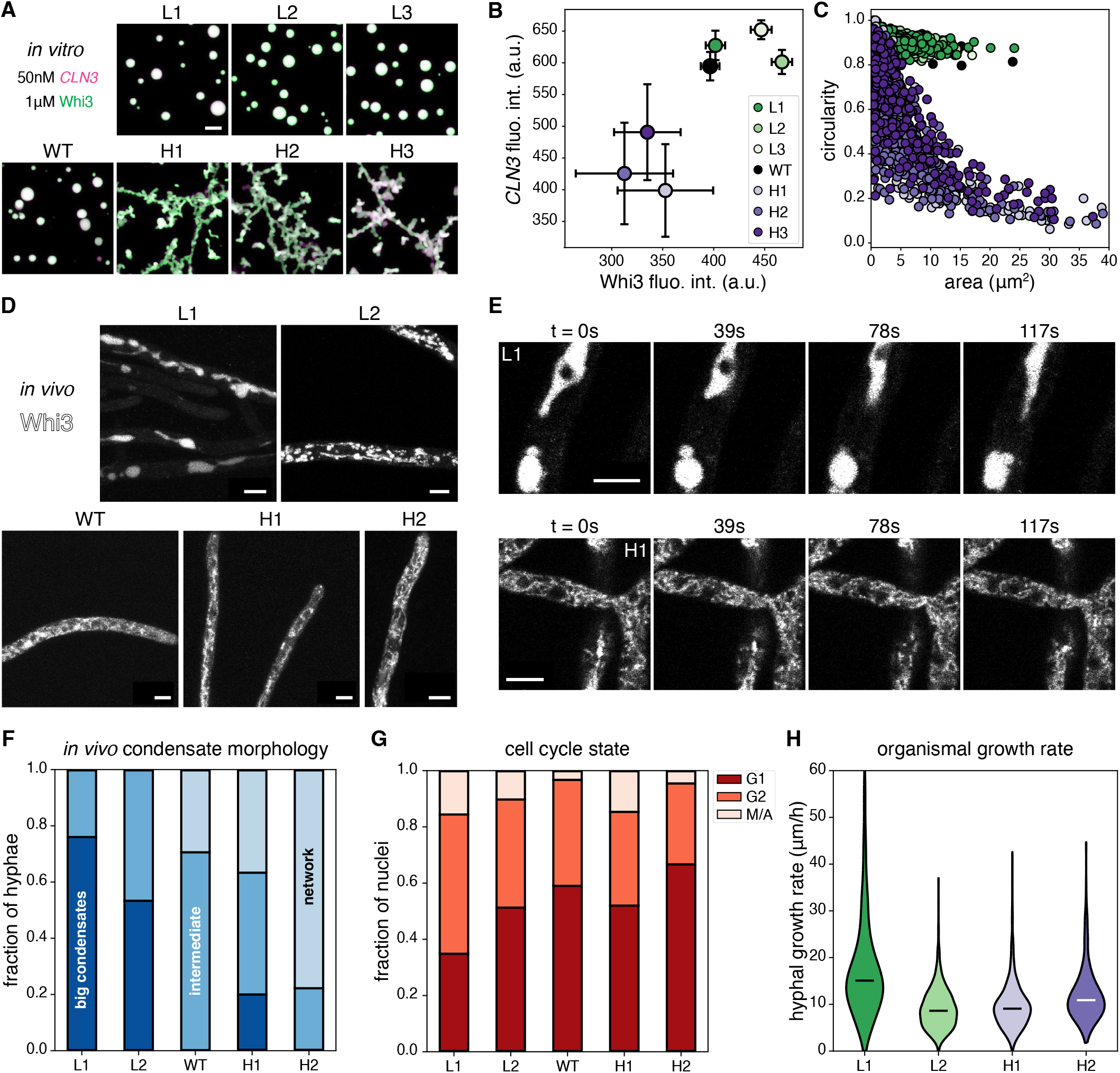
Upon phase separation, RNA ED encodes condensate material properties and can alter cell physiology. **(A)** Images are maximum z-projections of condensates formed by incubating 50nM of each *CLN3* structure mutant with 1μM Whi3 for 5 hours at 25°C. Green corresponds to Whi3 and magenta corresponds to *CLN3*. Each channel is contrasted identically in all images. All scale bars throughout this figure correspond to 5μm. **(B)** The central regions of the largest 50 condensates from experiments corresponding to panel (A) are used to estimate dense phase fluorescence intensities for Whi3 and *CLN3* (see **Methods** and **Fig S13**). Fluorescence intensities have been divided by 100. Error bars represent standard deviations. **(C)** The circularity is plotted against the area of each condensate from experiments corresponding to panel (A). **(D)** Images are maximum z-projections of Whi3-tdTomato in *Ashbya* strains with the indicated *CLN3* structural mutants integrated into the genome. Each image is contrasted separately to aid visualization. **(E)** Images corresponding to the indicated times from time lapses of Whi3 in the L1 (top) and H1 (bottom) *Ashbya* strains are shown.

Do the morphological differences encoded by these RNAs persist in the non-equilibrium context of live cells with RNA helicases and protein chaperones? To investigate this question, we integrated the L1, L2, H1, and H2 *CLN3* structure mutants at the endogenous locus and promoter as the only copy of *CLN3* into *Ashbya* cells also expressing Whi3-tdTomato (**Fig 3D)**. We found that the condensate morphologies seen *in vitro* were largely recapitulated *in vivo*. WT *CLN3* forms small condensates and some network-like condensates consistent with previous work and are likely on the ER based on morphology (*27*). Low ED *CLN3* mutants had either extremely large Whi3 condensates which were often spherical and resulted in very little signal in the dilute phase/cytosol (**Fig 3D**, L1 image), or a mixture of extremely large condensates and networks of intermediate-sized assemblies (**Fig 3D**, L2 image). In contrast, high ED *CLN3* mutants primarily exhibited networks of small, seemingly dynamically arrested condensates that filled the cytoplasm (**Fig 3D**, H2 and H1 images) and occasional larger puncta (**Fig 3F**). These mutant strains had comparable concentrations of *CLN3* RNA in the cytoplasm but some differences in degree of cell-to-cell variability of concentrations (**S14**). There was also some hypha-to-hypha variability in the appearance of condensates which we suspect reflects known differences in metabolism, growth, and cytoplasmic organization that are seen in these cells. Differences in material properties of Whi3 condensates in the low and high ED *CLN3* mutant *Ashbya* cells were apparent in timelapses. The L1 and L2 cells often showed large, flowing, liquid-like condensates (**Fig 3E**, top), while the H1 and H2 cells often showed relatively static and rigid, cytoplasm-spanning networks (**Fig 3E**, bottom) (**supplemental movies S1-S4**). Thus, RNA conformational heterogeneity similarly impacts properties of Whi3 condensates both *in vivo* and *in vitro* (**Fig 3A, D, F**).

We next assessed the functional consequences of different material states of Whi3-*CLN3* condensates. *CLN3* is a G1 cyclin and is responsible for progression through the cell cycle, so we measured the nuclear division state in the mutant cells using spindle pole body and nuclei staining (**Fig S15**). Previous work has shown that knockouts or mutations that eliminate Whi3-*CLN3* condensates in cells synchronize nuclear divisions (*28*). We found no significant differences in nuclear synchrony among the mutants indicating that the cell cycle is still regionally controlled despite different material states of Whi3 assemblies (**Table S3**) which may be expected since condensates persist in all strains. We did, however, observe that the L1 cells, which have the most pronounced increased droplet size phenotype, have a ∼40-50% reduction in the proportion of G1 nuclei and a ∼40-50% increase in the proportion of G2 nuclei, relative to the other mutants (**Fig 3G**). This observation suggests that the rate of progression through the cell cycle may be increased in the L1 cells which have exceptionally fluid-like and large condensates. We predicted this would lead to an increase in the nuclear density if the nuclei progress faster through the cell cycle, however, we surprisingly found no differences in the density of nuclei among the *CLN3* structure mutants (**Fig S16**). We therefore reasoned that the growth rate of L1 cells may be higher, and found indeed that the hyphal growth rate is ∼40-80% higher in the L1 cells than in the other strains (**Fig 3H**). These data suggest that the exceptionally large Whi3 condensates in the L1 cells can promote increased progression through the cell cycle and concomitant enhanced growth rates.

Circular black regions of exclusion correspond to nuclei. Each image is a single z-slice, and each row is contrasted separately to aid visualization. **(F)** Hyphae in images from experiments corresponding to panel (D) were categorized as belonging to 1 of 3 categories: “big condensates” as shown in the L1 image in panel (D), “intermediate” as shown in the L2 and WT images in panel (D), or “network” as shown in the H1 and H2 images in panel (D). The total numbers of categorized hyphae for each *CLN3* mutant *Ashbya* strain are L1 (75), L2 (30), WT (17), H1 (60), and H2 (18). **(G)** Spindle pole body and nuclei staining were performed on fixed *Ashbya* cells from which nuclear division states were determined (see **Fig S15**). The total numbers of categorized nuclei for each *CLN3* mutant *Ashbya* strain are L1 (109), L2 (39), WT (61), H1 (150), and H2 (66). **(H)** Hyphal growth rates were measured for each of the indicated *CLN3* mutant *Ashbya* strains. Black and white bars represent medians. The total numbers of measured hyphae are L1 (672), L2 (681), H1 (895), and H2 (675). All distributions are significantly based on the Mann-Whitney U test with p < 0.01, except for L2 and H1 which are different with p = 0.048.

## Opposing effects of RNA conformational heterogeneity on RNA clustering across the phase boundary

What is the molecular basis of the ED impacting the composition and form of condensates given the opposing trends seen above and below saturation concentrations? We investigated the surprising reversal of ED-dependent RNA recruitment across the phase boundary (**Fig 2B, Fig 3B**) by studying the nature of condensates formed just within the phase boundary using TIRF microscopy (**Fig 4A**). Relative to the subsaturated conditions shown in **Fig 2B**, we increased the Whi3 concentration by 50nM and observed small condensates for all *CLN3* structure mutant systems (**Fig 4A**). We segmented and quantified condensate properties and found that low ED *CLN3* systems form larger condensates (**Fig 4B**) that recruit more *CLN3* (**Fig 4C**) than their high ED counterparts. We conclude that low ED *CLN3* systems switch from recruiting less *CLN3* to recruiting more *CLN3* than high ED systems as they cross the phase boundary.

**Figure 4:**
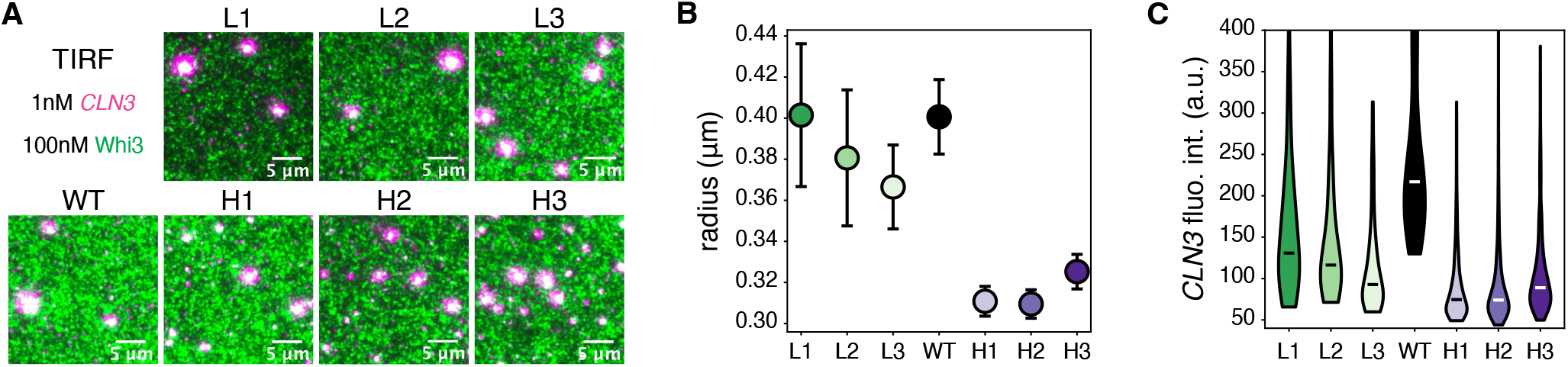
RNA conformational heterogeneity has opposite effects on RNA clustering across the phase boundary. **(A)** TIRF microscopy was used to visualize the first condensates formed as the phase boundary is crossed. The magenta channel corresponds to *CLN3* and the green channel corresponds to Whi3. Scale bars correspond to 5μm. **(B)** Condensates in experiments corresponding to panel (A) are segmented, and their radii are plotted for each *CLN3* structure mutant. Error bars represent 95% confidence intervals. **(C)** *CLN3* fluorescence intensity distributions are plotted for segmented condensates. Black and white bars indicate medians. All distributions are significantly different based on the Mann-Whitney U test with p < 0.01.

These data suggest that the phase boundary represents a change in the mechanisms by which RNA ED impacts condensate composition and properties. We hypothesize that RNA conformational entropy is a key piece of physical information encoded in the mRNA sequence itself (see supplementary text **Physical Information** and **Fig S17**). The ensemble diversity is a readily computable proxy for conformational entropy within a range of values which are relevant to our studies (**Fig S17B**). In subsaturated conditions, high conformational entropy exposes many single-stranded regions and increases the probability of productive RNA-RNA interactions (**Fig S17C**). However, we hypothesize that upon phase separation, there is a large entropic penalty associated with large-scale RNA networking within condensates, which limits partitioning for RNAs with high conformational entropy. Our hypothesis is based on the observation that high ED sequences tend to gain less or even lose ED upon dimerization (**Fig 2D**), and the prediction that this relationship becomes more extreme as RNAs form trimers, tetramers, and larger networks within condensates. For high ED sequences, we propose that the entropic penalty upon phase separation dominates the enthalpic gain of increased RNA-RNA interactions and explains the weakened recruitment of Whi3 and *CLN3* to condensates (**Fig 3B**). We propose that high ED sequences also lead to longer viscoelastic relaxation times due to enhanced networking within the dense phase, generating dynamically arrested assemblies *in vitro* and in cells (**Fig 3A, C, D, F**).

## RNA encodes physical information

These experiments show that RNA sequences can encode a hierarchy of information that includes both the genetic information to build a protein but also physical information that can impact the extent of RNA-RNA interactions in subsaturated assemblies and material properties of condensates. Because the genetic code is redundant with respect to amino acids specified by codons, there is extensive tunability in RNA conformational entropy and RNA-RNA interactions without altering the encoded amino acid sequence (**Fig 5**).

**Figure 5:**
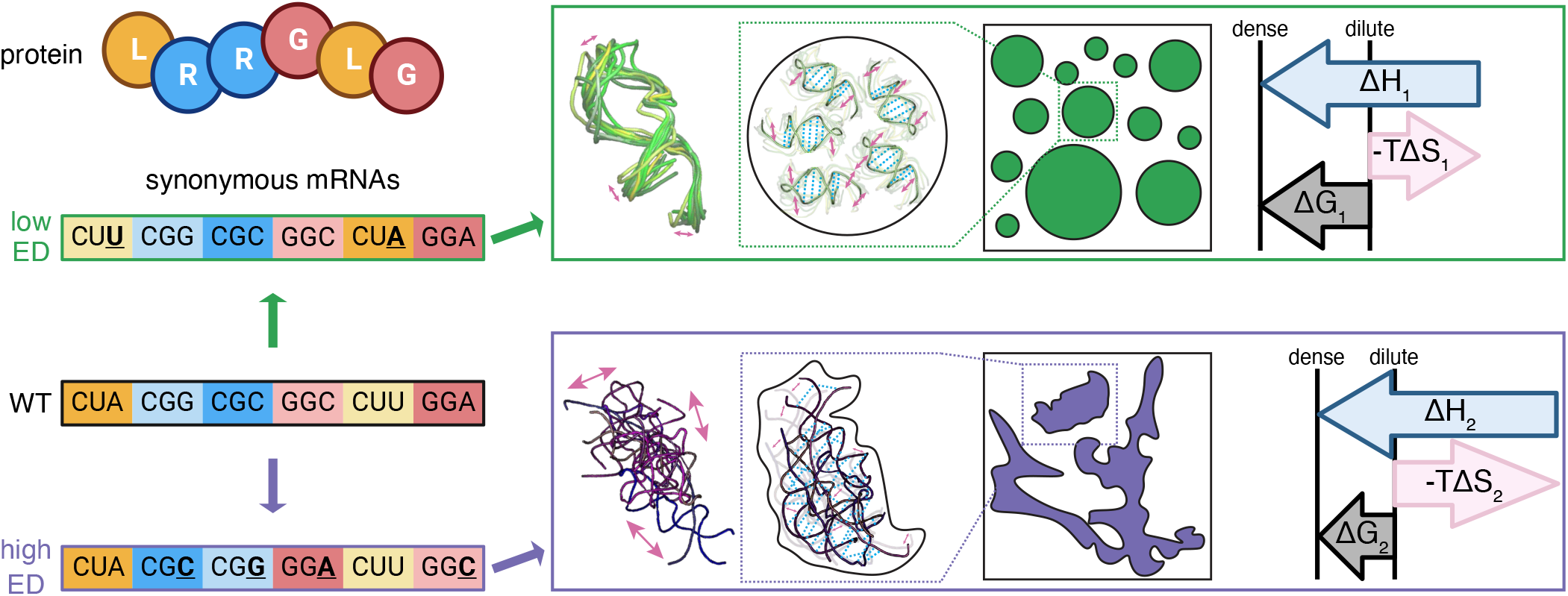
RNA encodes physical properties via disparate free energy costs of conformational entropy penalties. Synonymous mutations are designed and introduced to the WT mRNA to generate RNAs with polarized ensemble diversity. The low ensemble diversity RNA monomer samples a class of structures very similar to each other (top, green), which limits RNA-RNA interactions in the dense phase, but as a result, minimizes conformational entropy costs upon condensation. The high ensemble diversity RNA monomer samples a class of structures that greatly diverge from each other (bottom, purple), which lead to mesh-like RNA-RNA interactions in the dense phase and a high conformational entropy cost upon condensation.

We demonstrate that RNA can encode physical information, such as conformational heterogeneity, to influence condensate composition by balancing enthalpy contributions with conformational entropy costs (**Fig 5**). Similarly, proteins have long been recognized for this capacity to affect free energy landscapes through conformational entropy, impacting Boltzmann-weighted sampling across dense and dilute phases (*29, 30*).

RNA conformational heterogeneity may represent an orthogonal target of natural selection independent of protein sequence that nonetheless can impact the fate of the RNA’s localization, expression, or stability through condensate formation or smaller-scale assemblies sensitive to this property. It is clear that there are constraints on ED in the human genome (*31, 32*), potentially to maintain specific structures. However, examples of pathologies where synonymous mutations impact a pathology and are associated with altered ED exist, including in the critical oncoprotein KRAS (*32*). We predict that synonymous mutation might be a mechanism used by free-living organisms that can quickly adapt to rapidly fluctuating environmental conditions through tuning RNA conformations and condensate properties without altering protein-coding sequences. This study shows that RNA can encode physical information along with the message of the genetic code. We have demonstrated that this physical information can impart material properties to biomolecular condensates, which may drive different phenotypic outcomes. Similarly, the physical properties of RNA could play a significant role throughout its entire lifespan in the cell.

## Acknowledgments

We would like to thank the Gladfelter lab for many helpful discussions in the course of this study. We also would like to thank students in the Physiology course at the Marine Biological laboratory in Woods Hole, MA who piloted some of these experiments: Austin J. Graham and Arnold J.T.M. Mathijssen. We would like to thank Zhao Zhang from the Department of Pharmacology and Cancer Biology at Duke University for his contribution of Nanopore instrumentation GridIon.

## Author contributions

I.S., V.Z., A.P.J., B.M.S., S.J.C., and J.E. performed experiments. I.S. designed the RNA sequences. H.T.N. performed Langevin dynamics simulations. I.S., V.Z., A.P.J. B.M.S. and H.T.N. analyzed the data. I.S., V.Z., and A.S.G. drafted the paper, and all authors edited it.

## Data and materials availability

Data and materials are available upon request.

## Supplementary Materials

Materials and Methods

Supplementary Text

Figs. S1 to S18

Tables S1 to S3

Movies S1 to S4

## References

1. M. Vihinen, When a Synonymous Variant Is Nonsynonymous. Genes 13, 1485 (2022).

2. V. Mauno, Nonsynonymous Synonymous Variants Demand for a Paradigm Shift in Genetics. Current Genomics 24, 18–23 (2023).

3. F. Buhr et al., Synonymous Codons Direct Cotranslational Folding toward Different Protein Conformations. Molecular Cell 61, 341–351 (2016).

4. I. M. Walsh, M. A. Bowman, I. F. Soto Santarriaga, A. Rodriguez, P. L. Clark, Synonymous codon substitutions perturb cotranslational protein folding in vivo and impair cell fitness. Proceedings of the National Academy of Sciences 117, 3528–3534 (2020).

5. N. Sanchez de Groot et al., RNA structure drives interaction with proteins. Nature Communications 10, 3246 (2019).

6. K. L. Engel, A. Arora, R. Goering, H.-Y. G. Lo, J. M. Taliaferro, Mechanisms and consequences of subcellular RNA localization across diverse cell types. Traffic 21, 404–418 (2020).

7. K. A. Cottrell, R. J. Andrews, B. L. Bass, The competitive landscape of the dsRNA world. Molecular Cell 84, 107–119 (2024).

8. H. Wu et al., Unveiling RNA structure-mediated regulations of RNA stability in wheat. Nature Communications 15, 10042 (2024).

9. C. Roden, A. S. Gladfelter, RNA contributions to the form and function of biomolecular condensates. Nature Reviews Molecular Cell Biology 22, 183–195 (2021).

10. E. W. Martin, T. Mittag, Relationship of Sequence and Phase Separation in Protein Low-Complexity Regions. Biochemistry 57, 2478–2487 (2018).

11. J. M. Lotthammer, G. M. Ginell, D. Griffith, R. J. Emenecker, A. S. Holehouse, Direct prediction of intrinsically disordered protein conformational properties from sequence. Nature Methods 21, 465–476 (2024).

12. E. W. Martin et al., Sequence Determinants of the Conformational Properties of an Intrinsically Disordered Protein Prior to and upon Multisite Phosphorylation. Journal of the American Chemical Society 138, 15323–15335 (2016).

13. J. Ding et al., Visualizing RNA conformational and architectural heterogeneity in solution. Nature Communications 14, 714 (2023).

14. L. R. Ganser, M. L. Kelly, D. Herschlag, H. M. Al-Hashimi, The roles of structural dynamics in the cellular functions of RNAs. Nature Reviews Molecular Cell Biology 20, 474–489 (2019).

15. R. Lorenz et al., ViennaRNA Package 2.0. Algorithms for Molecular Biology 6, 26 (2011).

16. W. Ma, G. Zhen, W. Xie, C. Mayr, In vivo reconstitution finds multivalent RNA–RNA interactions as drivers of mesh-like condensates. eLife 10, e64252 (2021).

17. B. M. Stormo et al., Intrinsically disordered sequences can tune fungal growth and the cell cycle for specific temperatures. Current Biology 34, 3722–3734.e3727 (2024).

18. Z. Zhao, C. Jiang, Features of Recent Codon Evolution: A Comparative Polymorphism-Fixation Study. BioMed Research International 2010, 202918 (2010).

19. E. M. Langdon et al., mRNA structure determines specificity of a polyQ-driven phase separation. Science 360, 922–927 (2018).

20. H. Zhang et al., RNA Controls PolyQ Protein Phase Transitions. Molecular Cell 60, 220–230 (2015).

21. T. T. Bizuayehu et al., Long-read single-molecule RNA structure sequencing using nanopore. Nucleic Acids Research 50, e120–e120 (2022).

22. H. K. Wayment-Steele et al., RNA secondary structure packages evaluated and improved by high-throughput experiments. Nature Methods 19, 1234–1242 (2022).

23. H. T. Nguyen, N. Hori, D. Thirumalai, Condensates in RNA repeat sequences are heterogeneously organized and exhibit reptation dynamics. Nature Chemistry 14, 775–785 (2022).

24. M. Kar et al., Phase-separating RNA-binding proteins form heterogeneous distributions of clusters in subsaturated solutions. Proceedings of the National Academy of Sciences 119, e2202222119 (2022).

25. S. H. Bernhart et al., Partition function and base pairing probabilities of RNA heterodimers. Algorithms for Molecular Biology 1, 3 (2006).

26. I. Seim et al., Dilute phase oligomerization can oppose phase separation and modulate material properties of a ribonucleoprotein condensate. Proceedings of the National Academy of Sciences 119, e2120799119 (2022).

27. W. T. Snead et al., Membrane surfaces regulate assembly of ribonucleoprotein condensates. Nature Cell Biology 24, 461–470 (2022).

28. C. Lee, P. Occhipinti, A. S. Gladfelter, PolyQ-dependent RNA–protein assemblies control symmetry breaking. Journal of Cell Biology 208, 533–544 (2015).

29. Clifford P. Brangwynne, P. Tompa, Rohit V. Pappu, Polymer physics of intracellular phase transitions. Nature Physics 11, 899–904 (2015).

30. D. Scholl, A. A. Deniz, Conformational Freedom and Topological Confinement of Proteins in Biomolecular Condensates. Journal of Molecular Biology 434, 167348 (2022).

31. W. N. Moss, The ensemble diversity of non-coding RNA structure is lower than random sequence. Non-coding RNA Research 3, 100–107 (2018).

32. J. B. S. Gaither et al., Synonymous variants that disrupt messenger RNA structure are significantly constrained in the human population. GigaScience 10, (2021).

33. P. Eastman et al., OpenMM 7: Rapid development of high performance algorithms for molecular dynamics. PLOS Computational Biology 13, e1005659 (2017).

34. J. D. Honeycutt, D. Thirumalai, The nature of folded states of globular proteins. Biopolymers 32, 695–709 (1992).

35. E. Marinari, G. Parisi, Simulated Tempering: A New Monte Carlo Scheme. Europhysics Letters 19, 451 (1992).

36. M. R. Shirts, J. D. Chodera, Statistically optimal analysis of samples from multiple equilibrium states. The Journal of Chemical Physics 129, (2008).

37. B. Lee, F. M. Richards, The interpretation of protein structures: Estimation of static accessibility. Journal of Molecular Biology 55, 379–IN374 (1971).

38. M. S., FreeSASA: An open source C library for solvent accessible surface area calculations. F1000Research 5, (2016).

39. J.-Y. Tinevez et al., TrackMate: An open and extensible platform for single-particle tracking. Methods 115, 80–90 (2017).

40. D. Legland, I. Arganda-Carreras, P. Andrey, MorphoLibJ: integrated library and plugins for mathematical morphology with ImageJ. Bioinformatics 32, 3532–3534 (2016).

41. J. Wendland, Y. Ayad-Durieux, P. Knechtle, C. Rebischung, P. Philippsen, PCR-based gene targeting in the filamentous fungus Ashbya gossypii. Gene 242, 381–391 (2000).

42. A. S. Gladfelter, A. K. Hungerbuehler, P. Philippsen Asynchronous nuclear division cycles in multinucleated cells. Journal of Cell Biology 172, 347–362 (2006).

43. C. Lee, S. E. Roberts, A. S. Gladfelter, Quantitative spatial analysis of transcripts in multinucleate cells using single-molecule FISH. Methods 98, 124–133 (2016).

44. S. Berg et al., ilastik: interactive machine learning for (bio)image analysis. Nature Methods 16, 1226–1232 (2019).

45. J. R. Pringle, A. E. M. Adams, D. G. Drubin, B. K. Haarer, in Methods in Enzymology. (Academic Press, 1991), vol. 194, pp. 565–602.

46. S. E. R. Dundon et al., Clustered nuclei maintain autonomy and nucleocytoplasmic ratio control in a syncytium. Molecular Biology of the Cell 27, 2000–2007 (2016).

