## supplementary materials for "RNA encodes physical information"

**The PDF file includes:**

Materials and Methods
Supplementary Text
Figs. S1 to S18
Tables S1 to S3
Captions for Movies S1 to S4

**Other Supplementary Materials for this manuscript include the following:**

Movies S1 to S4

### Materials and Methods

Ensemble diversity

The ensemble diversity (ED) is defined as the Boltzmann-weighted average pairwise distance between all structures in the ensemble: $\text{ED} = p(S_i)p(S_j)d(S_i, S_j)$, where $S_i$ is the $i^{th}$ structure in the ensemble, $p(S_i) = \frac{1}{Q}\exp\left(\frac{-E(S_i)}{kT}\right)$ is the probability of structure $i$, where $Q = \sum_k \exp\left(\frac{-E(S_k)}{kT}\right)$ is the partition function, $E(S_i)$ is the energy of structure $i$, which is the free energy of folding as defined in Zuker's algorithm (*15*) and $d(S_i, S_j)$ is the distance between structures $i$ and $j$, defined as the number of base pairs needed to rearrange to transform structure $i$ into structure $j$. The ED can be equivalently defined in terms of the base pair probabilities, $p_{ij}$, which indicate the ensemble-level probability that nucleotide $i$ is bound to nucleotide $j$, as $\text{ED} = p_{ij}(1 - p_{ij})$. The units of ensemble diversity are nucleotides.

*in silico* RNA sequence design

We developed a genetic algorithm to design *CLN3* sequences with different structural properties. The algorithm is implemented in the 2 python scripts, 'evolve_CLN3_highED.py' and 'evolve_CLN3_lowED.py' that increase and decrease the ensemble diversity of the input *CLN3* sequence, respectively. Our algorithm relies on the RNAfold algorithm implemented in the python module RNA from the ViennaRNA software suite (*15*). The following parameters are set at the beginning of our algorithm: *fractional nucleotide content threshold*, which limits the change in the fractional content of each nucleotide relative to the input sequence (set to 0.01 for this study); *fractional codon content threshold*, which limits the change in the fractional content of each codon relative to the input sequence (set to 0.1 for this study); the *number of sequences to generate*, the *number of generations of mutants* (set to 1000 for low ED sequences and 10,000 for high ED sequences in this study); the *temperature* (set to 25C for this study); and the *salt concentration* (set to 150mM for this study). Next, the full WT *CLN3* sequence and just the *CLN3* coding sequence are input and the fractional nucleotide content, fractional codon content, amino acid sequence, and location of Whi3 binding sites (UCGAU) are all calculated. The first generation of mutants is next generated. The number of mutants in each generation corresponds to the number of available CPUs, since the structure evaluation is parallelized (128 for this study). In the first generation, each mutant is generated by choosing a random number between 5 and 50 from a uniform distribution which is the number of codons to mutate, and then randomly choosing the positions of the codons only within the coding sequence corresponding to the number of mutations, for each mutant. For each codon, a codon which encodes the same amino acid is randomly chosen with probability equal to its occurrence in the WT sequence. Once all mutants are generated, the free energy of their minimum free energy structure (MFE) is evaluated in parallel using the RNAfold algorithm. All sequences which violate the constraints of fractional nucleotide content, fractional codon content, and Whi3 binding site conservation are discarded. To generate high (low) ED *CLN3* mutants, the mutant with the least (most) negative MFE is chosen and becomes the parent of the next generation. In the following generations, if a mutant is generated with a more extreme MFE in the appropriate direction, it is automatically chosen as the next parent. However, if no such mutant exists, the most extreme candidate is chosen according to the Metropolis criteria in which the mutant is chosen with a probability given by $e^{-\Delta \text{MFE}}$, where $\Delta\text{MFE} = \left|\text{MFE}_{parent} - \text{MFE}_{mutant}\right|$. This allows for escape from local minima during the evolution process (**Fig S2A, B**). We find that MFE is tightly linearly correlated with ED (**Fig S2C**), and therefore use the MFE as the cost function since it is computationally simpler to calculate than the ED which relies of calculation of the full partition function. The following are the designed sequences used in the study:

WT *CLN3:*

gggagagucugcauaccaaagaucagccgcuugcauuaaaggggacgaaccggggcucuuauccuuagcaucgucucuuguccacaccucguugaccugcaga
cacagcacaaacccugcauaauuaauaguaaucauugucaucuccagcugggcuguuaauaucucauacccgaacaccuuugcauauuauacacauauucguc
auggacucuacaaaacucucaugcgauaagguuucgcugcuacaacuggccucaaucaggaggucuguguggggacaguaaggcaucccaucccccgguuaauac
acauggagaugguugcgcaccaaucugccugccaggaguacagcauugaaauccucaagcaucugauugaucucgagaaucagacccggucgucauggacuag
cuuccagucccagccggaacuuaccguggagaugcgcacuuugauuuucgacuuuaucaugagcugccacacucgcuugggacuuuccuccucgacacucuu
cuuaugcuacaacaucauugaucgguacugcuccaagaucauuguuaaaucacccacauaucagcuacuagggcuuaccgcucuuuggcuggcguccaaguuc
gcagacaaaaagccccgaauaccgagccugcagagccucugcugccaccagcacacgaaacagcaguucaaggaaauggagcugcacauccucaagagccuuaa
cuggucugugugcucugcuccgagucacgauucguucguggauauccuccugaaaaccaagauagccaaucucagcuucaagcgccucaaucuaaacgauuua
aaguauggagcaacgauucuuugcgagcuauccugcuuugacccagcacugaacuacaauuacaacuccagcgccauugccuuggcaagcguuacaguuauaa
ccugcgcacuccgacucgcugagcucuccgaguucauagauugcaggcgauguacuacugacaaaaaucucauugucaucugcaaccagcuucucugucuacu
agccugcgaggacaacuucccguccagcuuuaaccucaaguacgcuuccgagggcggcacgcuccacuccaaccccauuauggcccgucuguucgcuuauuac
cgccgguuagcugaugagaaaaaacguucguggcgugaugcaggcacucgcgugggcucgccaugugcaggcucagcaccggccagugcugccggcggcucc
cccgauauccgcauuggcggcgacacgcccccgggcucuguuccgccuuccgcucugggcccgcuugccccgcauaugccgacaaucaagacuuuugugccuc
ccacaccaacuacuccgagcaacucaucgcgcacucagcugucgcuuugugccaagccccauacgcuacccgcgccuguguccgucgcagcgucggcgcgcgca
aaaggcacugcacaucagcacacgccuccgguccagaaccccgcgcacuugcugcauacugcgcaaucuaagcgcgccgccucggccauggaccuugaguucu
ucgacauggaucaugcccuggucaagaaacuacgcuagcauac

L1 *CLN3*:

gggagagucugcauaccaaagaucagccgcuugcauuaaaggggacgaaccggggcucuuauccuuagcaucgucucuuguccacaccucguugaccugcaga
cacagcacaaacccugcauaauuaauaguaaucauugucaucuccagcugggcuguuaauaucucauacccgaacaccuuugcauauuauacacauauucguc
auggacagcacaaaguuguccugcgauaaggucagccucccuccagcuggccagcauaaggcgcucggugugggauucaaaggccagccaucccaggcuaauac
acauggagaugguugcccaccaaagcgccugccaggaguacucaaucgagauacuuaaacaucugauugaucuggaaaaccagacucgaagcuccuggacuag
cuuccagucccagccggagcugacaguggagaugcgaacucucauuuucgacuucauuaugagcugccacaccaggcuuggacuuaguucauccacgcuauuc
cuaugcuacaauaucaucgaucgauauuguagcaagaucauagugaagucaccaacuuaccagcuacugggccuaaccgcucucuggcuggccucaaaguucg
cggauaagaagccccggauccccucgcugcaaucgcuuugcugccaccagcacacuaagcagcaauucaaggagauggagcuccauauccucaaauccuugaac
uggucggugugcagugcucccucccacgacuccuuugucgauauuuugcucaaaacuaaaauagcaaaccugagcuucaaacgccuuaaccuuaacgaucuca
aguauggagcuacaauuuuaugcgagcuuaguuguuuugacccagcacuuaauuacaacuacaauuccagcgcuauugcguuggcuaguguuacugugauua
cgugcgcccugcggcucgcugaacucucugaguucaucgacugcaggcggugcaccaccgacaagaaccugauuguuauuugcaaucagcuucuuugucugu
uagcgugcgaggacaauuucccguccucauucaaucugaaguacgcuagcgagggcgggacucuccauucgaacccuaucauggcucgucuuuuugcuuauu
accgccgccucgcggaugagaaaaagcgcucguggcgcgacgcuggcacucguguggggucaccgugcgcaggcucagcaccggcaucugcugcaggcggca
gcccggauauccguaucggcggcgacacgccgccgggcucagugcccccucggcacugggcccgcuugcaccgcacaugccgacaauuaaaacuuucguucc
cccgacuccgacuacccccaguaacucgagucggacccagcugagccugugcgccaagccccacacgcugccagcaccuguguccguugcggcauccgcacggg
cuaagggcacugcgcaucagcacacacccccguucaaaaccccgcccaucuccugcauacugcgcagagcaaacgcgccgcgagcgccauggacuuagaguuu
uucgacauggaccacgcccuggucaagaaacuacgcuagcauac

L2 *CLN3*:

gggagagucugcauaccaaagaucagccgcuugcauuaaaggggacgaaccggggcucuuauccuuagcaucgucucuuguccacaccucguugaccugcaga
cacagcacaaacccugcauaauuaauaguaaucauugucaucuccagcugggcuguuaauaucucauacccgaacaccuuugcauauuauacacauauucguc
auggacagcacuaagcuguccugcgauaaaguguccuuauugcagcuugcaagcauccgccgcuccgugugggauucuaaggcgagccauccucgccuuauc
cacauggagaugguggcucaucagucagcgugccaggaguauucaaucgagauacucaagcaccugauugaucuugagaaucaaacccgcagcucauggacaa
guuuccaaucucaaccggaacuuacgguugagaugcgaacguugauuuucgacuuuaucaugagcugccauacgaggcucggccugagcagcuccacucuuu
uucucugcuacaacauaaucgaccgcuacuguagcaaaauuauuguuaagucgcccacuuaucagcuacucgggcucacugcccuguggcuagcaagcaaguu
ugcggauaaaaaaccccgcauuccuagcuugcagucccuaugcugccaucagcauaccaaacagcaguucaaggagauggagcugcacauucucaagagccuca
acuggucugugugcagcgcucccucccacgauucuuucguggacauccuacugaagacuaagauugcuaaucuuagcuucaagcgccuaaauuugaaugauc
ucaaauauggcgccacaauacuaugcgagcuuaguugcuucgacccagcgcugaacuacaacuacaacucuagugccauugcccucgcuuccgugacgguuau
uaccugcgcccuucgacuugcagaacucuccgaguucauagauugcaggcgaugcacgacagauaagaaucugauuguuaucugcaaucagcuucucugucu
gcucgccugcgaggacaacuuccccuccuccuucaaccugaaauacgccuccgagggaggcacacuccauaguaaccccauuauggcccgccucuucgcuuacu
accgccggcuggcugacgaaaaaaagcgcagcuggcgggacgcgggcacuagggucggcagcccgugugcaggcucagcaccagcgucagcugccggcggcuc
uccggacauccggaucggcggcgacacuccgccaggcagugucccgccgagugcacugggcccgcuggccccgcacaugccgaccaucaagacuuuuguccccc
ccgacacccacuacucccucgaacucuucgaggacgcagcugucgcuuugugcuaagccgcacacgcugccggccccggucucgguugcggcaagcgcgcgug
ccaagggcacugcacaccagcacacuccgccugugcagaaccccgcgcacuugcugcauaccgcacagucgaaacgugcagccucugcaauggaccuagaguuc
uucgauauggaucacgcgcuggucaagaaacuacgcuagcauac

L3 *CLN3*:

gggagagucugcauaccaaagaucagccgcuugcauuaaaggggacgaaccggggcucuuauccuuagcaucgucucuuguccacaccucguugaccugcaga
cacagcacaaacccugcauaauuaauaguaaucauugucaucuccagcugggcuguuaauaucucauacccgaacaccuuugcauauuauacacauauucguc
auggauuccacgaagcucagcugugauaaaguuucccuccuucagcuggcaaguaucaggaggagcgugugggacucgaaagccagccaucccc gucucauuc
acauggagaugguugcccaucagagugcuugccaggaguacucuaucgagauccucaagcaccugauugaucucgaaaaucaaacccggucgagcuggaccuc
uuuccagucucagccggaacuaaccguggagaugcgcacuuugauuuucgauuucauuaugagcugucacacucggcucgggcuuucgaguuccacucucuu
ccugugcuacaacauuauugaccgguacugcagcaagaucauaguuaagagccccaccuaucaacuccuaggccugacugcuuuguggcucgcgagcaaauuc
gcggacaaaaaaccgcgaauucccucgcugcagagucucugcugccaccagcacacaaagcagcaguucaaggagauggagcugcacauccucaagucucuuaa
cuggagcgugugcucagcuccaucccacgauuccuucguggacauccuucuaaaaacgaagaucgcgaaccucagcuucaaacgccuaaauuuaaacgaucug
aaguauggagccacgauacuuugugagcucucaugcuucgauccggcucuuaacuacaacuacaacucuagcgcaaucgcccuagcaucuguuacggugauaa
cuugcgcacuccgccucgccgagcuaucggaguucauagauugccgccguugcacaacugacaagaaucuuauugucauuugcaaccagcuucuaugccuccu
cgcaugcgaggacaauuucccguccucguuuaaccucaaauacgcaucggagggcgggacucuacauagcaaucccaucauggccaggcucuucgccuauuac
cggcgccuggccgaugagaaaaagcgcucguggcgagaugcaggcacccgcgugggcucgccaugugcgggcucagcuccggccagugcggccggcggcucu
cccgauauccgaauaggcggcgacacgccgcccggaucggugccgccguccgcacuuggcccgcuggccccgcacaugccgacaauuaaaacuuucguuccgc
ccacaccaacaacuccguccaacagcucuaggacgcaguugucccugugugccaaaccucacacacugcccgcgccugugucuguugccgcgagcgcucgggc
caagggcacugcccaucagcauacgcccccgguccagaaucccgcccacuugcugcauaccgcucaguccaagcgggcugcguccgcuauggaccuggaguuu
uuugacauggaucacgcucuggucaagaaacuacgcuagcauac

H1 *CLN3*:

gggagagucugcauaccaaagaucagccgcuugcauuaaaggggacgaaccggggcucuuauccuuagcaucgucucuuguccacaccucguugaccugcaga
cacagcacaaacccugcauaauuaauaguaaucauugucaucuccagcugggcuguuaauaucucauacccgaacaccuuugcauauuauacacauauucguc
auggauuccacgaaauuaagcugcgauaaagugucacugcuacaacuggccaguauccguagguccguuugggauaguaaggcaucgcauccgcgucugauu
cauauggagaugguggcccaccaaagcgccugucaagaauauagcauugagauacugaagcaccugaucgaccucgagaaccagacccguaguucguggaccu
cguuccagucgcagccggagcuuacuguggagaugcggacguuaauauuugauuucauuaugaguugccacacacgccucggccuaucuucgucuacuuuau
uccuuugcuacaacaucauugaccgguacugcuccaagauuauuguuaagucgccuacauaccaguuacugggccuuacagcucucuggcucgccuccaaguu
cgccgacaaaaagccucgcauaccuucacuccagucccuauguugccaccagcacacgaaacagcaguucaaagaaauggaauuacacauccuaaagucacucaa
uugguccguuugcuccgccccgucgcaugauucauucguugauauucuguuaaaaacaaagauugccaaccuuagcuuuaaacgccucaaccugaacgaccua
aaauauggcgcuacgauacuuugcgaguugaguugcuucgacccagcguuaaacuacaauuacaauuccuccgcaaucgcgcucgccucugucacugucauca
cuugcgcccuacgucuugccgagcuuuccgaguuuauagauugccgccgcugcacuacugauaagaacuugaucgugaucugcaaccagcugcuuugccuac
ucgcaugugaagauaauuucccgucaagcuuuaaccuuaaguacgcuuccgagggggcacacugcacuccaacccaauuauggcacgccuauucgcauauua
ccgacgccuggccgaugagaagaagaggucauggagggacgcaggcacacgcguuggcucuccgugcgcaggaucggcccccgcaagcgcugccggaggcuc
gcccgauauucgcauuggcggagacaccccgcccggcucuguuccaccuuccgcacuuggcccgcucgccccgcacaugccuacuaucaagacuuucgugccg
cccacaccgacuacccccuccaauucgucccgcacucaguugucgcuuugcgcaaagccccauacccucccccgcgcccguguccgucgcagcauccgcacgagc
uaaaggcaccgcacaccagcauacaccccccguccagaaccccgcucaucuacugcauacugcccagucuaagcgagccgccuccgccauggaucucgaguucu
ucgacauggaucacgcacuggucaagaaacuacgcuagcauac

H2 *CLN3*:

gggagagucugcauaccaaagaucagccgcuugcauuaaaggggacgaaccggggcucuuauccuuagcaucgucucuuguccacaccucguugaccugcaga
cacagcacaaacccugcauaauuaauaguaaucauugucaucuccagcugggcuguuaauaucucauacccgaacaccuuugcauauuauacacauauucguc
auggacagcacuaaguugaguugcgauaaaguuagccuguuacagcuugccucgauccgccgcagcgugugggauuccaaagcuucgcauccccggcucauu
cauauggagaugguggcccaccagucagccugccaagaguacaguauugagauccuaaagcaucugaucgacuuagagaaccagacacgaaguucguggacgu
cauuccaaagccagccggagcuuaccguggagaugcgcacucucauauuugauuucaucaugucuugccacacccgccucggacuauccuccucgacacuuuu
ccucugcuacaauauaauugauagguacugcucgaagauuaucgugaagucgccgacguaucagcuccuuggccuaacagcacuaugguuagcaucgaaguu
ugccgacaaaaagccgcggauucccagccuucagagccuguguugccaccagcauacuaagcagcaauucaaggagauggagcuccacauucuaaaaucccuca
acugguccguguguuccgcccccucucaugacucuuucguugauauccuccugaaaaccaaaauugcgaaccugagcuucaaacgccugaaucuaaaugaccu
aaaguauggcgcuacuauuuuaugcgagcuauccuguuucgacccggcucuaaacuauaacuacaauucaucagcgaucgcccuggcuuccgucacuguuaua
acuugcgcucugcgucuugccgaacugucugaauucauugacugccgccgauguacaacugauaagaaccugaucgugauauguaaccagcuccuaugccuac
ucgcuugcgaggauaacuucccgucgucguucaaccucaaauacgcuucagaaggcggcacucuccacagcaauccgauuauggcacgccucuucgcauacua
ccgccgucuggccgacgagaagaaaaggucuuggcgugacgcaggaacucgcgugggcaguccgugcgccggcagcgccccagccucagccgcaggcggcucu
cccgauauacgaauuggcggcgauacucccccuggcuccguuccuccuuccgcacuuggcccgcucgcaccccacaugccgaccauaaagacuuucguuccgc
cuacgccuacuaccccaucuaauagcucccgaacgcagcucucauuaugcgcaaagccgcacacacuacccgcccccgucucagucgccgcguccgcacgcgca
aaaggcacggcacaccagcacacaccaccgguucagaaccccgcacaucuucugcauacagcacagucgaaacgcgccgcuuccgccauggaucuagaauucuu
ugauauggaucacgcucuggucaagaaacuacgcuagcauac

H3 *CLN3*:

gggagagucugcauaccaaagaucagccgcuugcauuaaaggggacgaaccggggcucuuauccuuagcaucgucucuuguccacaccucguugaccugcaga
cacagcacaaacccugcauaauuaauaguaaucauugucaucuccagcugggcuguuaauaucucauacccgaacaccuuugcauauuauacacauauucguc
auggacagcacuaaauugaguugcgauaaggucagccuccugcagcuagcauccauucgucguucaguuugggauucgaaagcaagccacccccgccuaauac

acauggagauggucgcccaccaguccgcaugccaagaguauagcauugagauacuaaagcaccuuaucgaccuagagaaucagacccgcucgaguuggacgag
uuuccaaagccaaccggaacuaacuguggagaugcggacgcuaauuuucgauuucauaaugucgugccacacucgccuaggccugucgucguccacgcuauuc
cucugcuacaacaucauugaccgguauugcucgaagauaaucgugaaguccccuacguaccagcuacuuggccuuacugcgcuuuggcucgcaagcaaguucg
ccgauaaaaagccgaggaucccgucccuucagucgcuuuguugucaccagcacacuaaacagcaguucaaagagauggagcuacauauccuuaagucccugaa
uuggucgguuuguucagccccgucccaugauaguuuuguugauauuuugcucaagacgaagauugccaaccuguccuucaagcgccucaaccucaacgaccu
caaguacggagccacgauucuaugugaacuuucaugcuucgacccggcccuuaacuacaauuacaauagcuccgcaauugcacucgccucuguuaccgugauc
accugcgcccuucgcuuagccgagcucuccgaguuuauagacugccgccgauguacaacugauaagaaccuaauugugauaugcaaccagcuguugugcuuac
ucgccugcgaagauaauuucccgucaucauucaaucugaaguacgcguccgagggcggaacucuucacuccaaccccaucauggcucgccuauucgcauacua
ucgacguuuagccgaugagaaaaaacgaagcuggcgugacgcaggcacucguguuggcuccccgugcgcaggcagcgcccccgcuuccgccgcuggcggcuc
uccagacauccgcauuggaggcgauacacccccuggcucggugccgcccucagcucuuggcccgcucgcuccucauaugccgacuauaaagacguucgugccg
ccuacuccgacuacacccuccaacuccucucgcacucaguuaucguuaugcgccaaaccucacacacugcccgcccccguguccgucgcagcaucagcacgcgc
uaaaggcaccgcccaucagcacacacccccguccagaaccccgcccaccuccugcauaccgcccaguccaaacgcgccgcaucugcaauggauuuagaguucu
uugauauggaucaugcgcuggucaagaaacuacgcuagcauac

<u>Sequence alignment of *CLN3* structure mutants</u>

Pairwise sequence alignments were performed among the seven RNA sequences (one wild-type and six mutants) using CLUSTALW. The algorithm employs the Needleman-Wunsch method for global alignment and calculates sequence identities to quantify conservation across the sequences.

<u>*in vitro* transcription</u>

The three high ED and the three low ED *in silico* designed RNA sequences were each synthesized as a DNA gene block by Twist Bioscience (San Fransico, CA). We then directly cloned each gene block into the pJET plasmid using the CloneJet (ThermoFisher, #K1231) kit according to the manufacturer's instructions. Each plasmid was sanger sequenced in the forward and reverse direction using the manufacture supplied primers (fwd: CGACTCACTATAGGGAGAGCGGC, rev: AAGAACATCGATTTTCCATGGCAG) by Azenta Life Science (South Plainfield, NJ).

To generate template for *in vitro* RNA transcription we PCR amplified each *CLN3* sequence with an upstream T7 using PCR (fwd: TAATACGACTCACTATAGGGAGAGTCTGCATACCAAAGATCAGCCGCTTGC, rev GTATGCTAGCGTAGTTTCTTGACC). We verified that the resulting fragment was the correct length and then removed any remaining plasmid DNA by digestion with DpnI (New England Biolabs, R0176).

*CLN3* RNA was synthesized using the Hi-Scribe T7 kit (New England Biolabs E2040) according to the manufacturer's instructions. Briefly, 1ug of DNA template was added to a tube with 2µl of 10x reaction buffer, 2µl of Hi-T7 polymerase, 0.5ul of RNAse inhibitor, 2µl of each NTP at 100mM, 0.1µl of Cy-5 labeled UTP and enough water to bring the total volume to 20µl. RNA synthesis was performed on a thermocycler set to incubate at 37°C for 4hr. Following synthesis, the solution was diluted to 50µl with RNase-free water and the DNA was digested by addition of 2µl of RNAse-free DNAse directly to the tube (New England Biolabs, M0303). RNA was precipitated by addition of 25µl of 2.5M LiCl followed by chilling at -80°C for 30min. The RNA was pelleted by centrifugation at maximum speed (20,000rcf) for 10min, washed briefly with 70% Ethanol and then resuspended in nuclease free water. The concentration was determined by A260 measuring using a Nanodrop. Purity was assessed by denaturing gel electrophoresis.

The label % of each purified RNA was calculated by measuring total RNA concentration and total Cy-5 concentration using a Nanodrop (Thermo Fisher). The ratio of the dye concentration to the RNA concentration gives the label %. The label % for the RNAs used in this study are: L1 – 46%, L2 – 46%, L3 – 46%, WT – 48%, H1 – 50%, H2 – 46%, H3 – 47%.

Direct-RNA Nanopore Chemical Probing

***in vitro* transcription**: Since Nanopore sequencing reads from 3' to 5' and the first few nucleotides are often not recorded at high confidence, we installed a 10 nucleotide extension at the 3' end of each RNA (**Table S2**). RNAfold simulations were used to ensure that the predicted ED of each RNA did not vary more than 2% relative to the predicted ED of the sequence without the 10 nt. extension. To avoid the effect of chemical probing in the extension, we use only G, C, and U nucleotides in the extension sequences. HiScribe T7 High Yield RNA Synthesis Kit was used to synthesize RNA using *in vitro* transcription (IVT). After incubation, the RNA product was precipitated using lithium chloride and resolvated in RNase-free water.

**Chemical probing of RNA**: All structural probing was carried out using Diethyl pyrocarbonate (DEPC) 96% (Sigma), which carbethoxylates unpaired adenosine (A) nucleotides at N-6 or N-7 by opening the imidazole ring. Single-stranded adenosines are reported by NMR to probe at 24.7% ± 4.27% (*21*). For each *CLN3* structure mutant, 10 µL of 500 nM RNA was denatured at 90ºC for 3 minutes, cooled on ice and renatured in 150 mM KCl, 50 mM HEPES pH 7.4 (phase separation buffer) for 20 minutes at 20ºC. For probing experiments with DEPC, RNA was treated with 10% DEPC for 45 minutes at 20ºC. The reactions were immediately purified using RNAClean XP beads (Beckman Coulter).

**Library preparation**: Sequencing libraries were prepared following the direct RNA sequencing protocol RNA-004 (Oxford Nanopore Technologies, ONT). The reverse transcription adapter (RTA) was replaced with sequence specific 3' adapters which are composed of oligo A and oligo B (**Table S2**). Oligo A is the same for all sequences and oligo B is RNA-specific. All oligos were ordered from IDT. RNAs are 3' adapted by mixing with T4 DNA ligase and 3' adapter duplex oligo which is made by mixing oligo A and pooled oligo B at a 1:7 ratio at 1.4 µM in annealing buffer (10 mM Tris-HCl pH 7.5, 50 mM NaCl), heating to 95ºC for 2 minutes, and cooling down slowly (0.1ºC / second). After renaturation and treatment with DEPC, 3' adapter-ligated RNAs were reverse transcribed using superscript III (NEB). The reverse transcription step serves to prop up RNA using its complementary cDNA to obtain longer reads and minimize the impact of RNA structure on nanopore reading. Note that the direct-RNA sequencing method sequences the RNA through Nanopore, not the cDNA.

**Direct-RNA Nanopore sequencing and alignment**: The reverse transcribed RNAs were pooled for each experimental condition and a motor protein ligated, which were subjected to Oxford Nanopore sequencing using MinION Flow Cell (RNA) on GridION Mk1 for 72 hours. Reads were basecalled using Super accurate (SUP) model (Oxford Nanopore Technologies, ONT). Reads were aligned using ONT software to target RNAs allowing for mismatches, insertions, and deletions to account for nanopore sequencing errors as well as probing adduct. Since all RNAs share less than 82% sequence identity, no demultiplexing was necessary to parse RNA reads into correct references.

**Calculation of pairwise distance**: Each read describes a portion of one conformation within the RNA structural ensemble, as reads do not encompass the entire sequence. To analyze this data, we calculated the pairwise distances among the 1000 reads with the highest coverage for a given sequence (resulting in 1,000,000 read pairs), using the following equation:

$$d = \frac{N_D}{C}$$

Where $d$ is the pairwise distance between each pair of reads, $N_D$ is the number of adenosine discrepancies (probed in one read but not the other), and $C$ is the number of overlapping adenosines, or coverage, between the two reads. Note that $\langle d \rangle$ is proportional to the distance metric which is used in the calculation of the ensemble diversity, i.e. the number of base pairs needed to open or close to transform one structure into another, except that it only accounts for adenosines and is normalized by the read coverage to account for differences in read length and quality. A schematic of the analysis is shown in **Fig S4**.

Jensen-Shannon distance calculation

For the calculation of the Jensen-Shannon distances (JSD) shown in **Figs S5, S10** and corresponding to the data in **Figs 1D, 2B**, the following procedure was used. The maximum values among all distributions to be compared was computed, and probability vectors for all distributions were computed in 1000 linearly spaced bins ranging from 0 to this maximum value. For each pair of distributions, the JSD was calculated using the corresponding probability vectors using the function jensenshannon from the python library scipy.spatial. These JSDs are shown in the heatmaps in **Figs S5, S10**. The average JSD values among the low ED mutants average the 3 pairwise JSDs among L1, L2, and L3, and similarly for the average JSD values among the high ED mutants. The average JSD values among the high and low ED mutants average the 9 pairwise JSDs that arise from comparing L1, L2, and L3 to each of H1, H2, and H3.

Langevin dynamics simulations

We performed simulations using an updated version of the Single Interaction Site (SIS) (*23*). The modification restricts the number of multiple base pairs a single nucleotide can form simultaneously. Simulations were performed on Graphics Processing Units (GPUs) using OpenMM code (*33*) to enhance sampling of the conformational space. We used low friction Langevin dynamics, in which the viscosity of water was reduced by a factor of 100 (*34*). The simulations are computationally extensive due to the large RNA sizes, thus we used the simulated tempering method to ensure that the conformational space is sampled exhaustively (*35*). The trajectories were analyzed using the Multistate Bennett Acceptance Ratio (MBAR) to calculate all properties of interest (*36*). All simulations were performed in 1M NaCl, where electrostatic interactions are weak, and only base pairing interactions dominate. Simulations were performed for a single RNA molecule in the extended conformation 5 x $10^9$ time steps, in which the first 5 x $10^8$ steps were discarded, which ensures that only equilibrated structures are used in computing all quantities of interest.

Base accessibility of the nucleotide $i$ is defined as $H_i = S_i\gamma_i$, where $S_i$ is the solvent-accessible surface area of nucleotide $i$ (calculated using the Lee-Richards algorithm (*37*)) in FreeSASA (*38*). $\gamma_i$ adopts two values: 0-if the nucleotide i is involved in base pairing and 1-otherwise. In addition to the RNA secondary structure (reflected in $\gamma_i$), the accessibility of a base also depends on the RNA tertiary structure (reflected in $S_i$). If a base is deeply buried in the core of the RNA, its accessibility is small. On the other hand, if the base is located near the RNA periphery, its accessibility is high.

Native RNA gels

6µL of 2 µM of each RNA in 150 mM KCl 50 mM HEPES pH 7.4 (phase separation buffer) was loaded into a 1% agarose gel in TBE (89 mM Tris-HCl pH 7.8, 89 mM borate, 2 mM EDTA) with a 2:50000 volume fraction of SYBR Safe DNA gel stain (Thermo Fisher) and imaged with UV.

Whi3 protein purification

Whi3 constructs were transformed into BL21 competent cells (New England Biolab C2527I). Cells were grown at 37C with shaking and appropriate selection until an OD600 of 0.6 was reached. Then cells were shifted to 18C and induced with 1mM IPTG (Fisher I2481C25). Cells were grown for 18hr and then pelleted by centrifugation at 13000 rcf. The Pellet was resuspended in 25ml

Lysis buffer (1.5M KCl, 20mM Imidazole, 5mM beta-mercaptoethanol, 50mM Hepes pH7.4) along with 1 tablet Pierce Protease inhibitor (Thermo-Fisher Scientific A32965) and 15mg Lysozyme (Fisher BP535-10). Solution was transferred to a dounce homogenizer and processed until homogenous. This solution was transferred to a glass beaker on ice and sonicated at 70% power for 1 minute on, 2 minutes off, for 4 cycles. Lysate was clarified by centrifugation at 50000 rcf for 30min. Clarified lysate was mixed with 0.8mL of Cobalt resin (Thermo Fisher Scientific PI89965), column was washed with 5 column volumes of lysis buffer and then eluted with 3ml of elution buffer (150mM KCl, 200mM Imidazole, 5mM beta-mercaptoethanol, 50mM Hepes pH7.4). Protein was dialyzed into storage buffer (150mM KCl, 50mM Hepes pH 7.4) using a slide-a-lyzer cassette and a 20,000MW cutoff (Thermo-Fisher Scientific 87735). Protein was labelled using an atto-488-NHS ester (Sigma Aldrich 41698) and then dialyzed again to remove unconjugated dye.

Total internal reflection fluorescence (TIRF) microscopy

For the experiments corresponding to **Fig 2A, B**, 50nM Whi3 with a very low label % (<0.1%) was mixed with 1nM of each cy5-labeled RNA in 150 mM KCl 50 mM HEPES pH 7.4 (phase separation buffer) and incubated in a low-bind tube for 1 hour at 25ºC, pipetted onto plasma-cleaned glass, and imaged with TIRF microscopy. For the experiments with RNA only, 500pM of each RNA was mixed in phase separation buffer, pipetted onto plasma-cleaned glass, and imaged with TIRF microscopy. Spots in images were detected using the FIJI plugin TrackMate (*39*) and the average intensity of the *CLN3* fluorescence was extracted from each detected spot.

For the experiments corresponding to **Fig 4**, 100nM Whi3 with a very low label % (<0.1%) was mixed with 1nM of each cy5-labeled RNA in 150 mM KCl 50 mM HEPES pH 7.4 (phase separation buffer) and incubated in a low-bind tube for 1 hour at 25ºC, pipetted onto plasma-cleaned glass, and imaged with TIRF microscopy. Since larger condensates were formed in this concentration regime, we used a different image analysis pipeline that uses custom FIJI macros and python scripts. The first script 'TIRF_analysis_sep2023-STEP1.ijm' sets an automatic Otsu threshold in the RNA channel, runs the watershed operation, extracts areas and centroid, and saves them both. The python script 'images_from_centroids.py' creates a binary image corresponding to the centroids. A second FIJI macro 'TIRF_analysis_sep2023-STEP2.ijm' opens these centroid images, uses MorphoLibJ (*40*) to create ROIs around the centroids which are disks with radius 2 pixels, multiplies these ROIs by the original images to select only the dense phase signal in the center of each droplet, sets an Otsu threshold, and extracts mean intensities for each droplet. The script 'process_output2_TIRF.py' analyzes and plots the size and intensity data which is shown in **Fig 4B, C**.

RNAcofold predictions

We generated predictions about the structural ensembles of homodimers using the function RNAcofold from the python module RNA corresponding to the Vienna RNA software suite (*25*). For each RNA evaluated in **Fig 2C, D**, we calculated the free energy of the structural ensemble for the monomer, $G_{\mathrm{mono}} = -RT \ln(Q_{\mathrm{mono}})$, where $R$ is the gas constant, $T$ is set to 25C, and $Q_{\mathrm{mono}}$ is the partition function for the monomer, and the salt is set to 150mM; the ensemble diversity of the monomer, $\mathrm{ED}_{\mathrm{mono}}$; the free energy of the structural ensemble for the homodimer $G_{\mathrm{dim}}$; and the ensemble diversity of the homodimer, $\mathrm{ED}_{\mathrm{dim}}$. We calculated dimer $\Delta G = G_{\mathrm{dim}} - 2 \times G_{\mathrm{mono}}$ and dimer $\Delta\mathrm{ED} = \mathrm{ED}_{\mathrm{dim}} - 2 \times \mathrm{ED}_{\mathrm{mono}}$ which are the y-axis quantities shown in **Fig 2C, D**.

*in vitro* phase separation assays

**Preparation and microscopy:** Whi3 and *CLN3* aliquots were thawed on ice just before experiments. Whi3 aliquots were spun down at 13,200 rpm for 5 minutes to remove any aggregates, and Whi3 and its Atto 488 label % were calculated for each aliquot. Labeled and unlabeled Whi3 and labeled *CLN3* were diluted into buffer containing 50 mM Hepes at pH 7.4 and 150 mM KCl to desired concentrations and labeled fractions of protein and mRNA. The volumes of labeled and unlabeled protein stock needed to simultaneously obtain a desired total protein concentration and labeled fraction were determined using the following formulas: $u = \frac{VC}{[u]+\lambda[l]}, l = \lambda u, \lambda = \frac{L[u]}{(l_L - L)[l]}$, where *u* is the desired volume of unlabeled protein (microliters), *l* is the desired volume of labeled protein (microliters), *V* is the final reaction volume (200 µL for phase separation assays) *C* is the desired final protein concentration (micromolar), $[u]$ is the measured concentration of the unlabeled protein stock (micromolar), $[l]$ is the measured concentration of the labeled protein stock (micromolar), $l_L$ is the measured labeled fraction of the labeled protein stock, and *L* is the desired protein labeled fraction (5% - 10% for phase separation assays). Once mixed, 200µL solutions of Whi3 and *CLN3* were loaded into glass-bottom imaging chambers (Grace Bio-Labs) that were pre-coated with 30 mg/mL BSA (Sigma) for 30 minutes to prevent protein adsorption to the glass and sides of the well. The wells were washed thoroughly with buffer before adding the Whi3 and *CLN3* solution. Final solutions were mixed by pipetting without introducing bubbles, and incubated at 25C for 5 hours before imaging. Confocal z-stacks with 7 slices at 1µm intervals were acquired using both 488nm and 561nm lasers in 5 positions in each imaging well.

**Analysis:** Images are analyzed using custom FIJI macros and python scripts. The first script 'blob_analysis_july2024-STEP1.ijm' does the following: For a given z-stack, each z-slice is automatically thresholded in the Whi3 channel using the Moments dark threshold, and the z-slice which results in the largest summed area across all detected objects is selected for further analysis. The watershed operation is performed. Analyze particles is run and records the areas and centroids of each droplet. The python script 'images_from_centroids.py' is run next. It creates a binary image corresponding to the centroids. A second FIJI macro 'blob_analysis_july2024-STEP2.ijm' opens this binary file, uses MorphoLibJ to create ROIs corresponding to the centroids by growing disks with radius 2 pixels, multiplies these ROIs by the original z-slice in each channel to select out only these central regions of each droplet, sets an Otsu threshold, and runs Analyze particles to extract mean intensities for each channel and each region. The output data is further processed using the script 'phase_diagrams.py' to produce phase diagrams shown in **Fig 3B, S12**. The fluorescence intensities of *CLN3* and Whi3 are normalized by their label %. The intensities corresponding to the 50 droplets with the largest radii in each condition are selected, since the dense phase intensities no longer vary with size in this range (**Fig S13**). These values are used to compute means and standard deviations of the dense phase fluorescence intensities shown in the phase diagrams. For the analysis of circularity vs area shown in **Fig 3C**, a different FIJI macro was used. 'blob_analysis_morphology_july2024.ijm' uses only the protein channel, and selects the z-slice corresponding to the largest summed area as before using the Moments dark threshold. The watershed operation is not used, however, and the area and circularity of the detected objects in the chosen z-slice are recorded. The resulting values are processed and plotted with the script 'analyze_morphology.py'.

Integration of *CLN3* structure mutants in *Ashbya*

To generate endogenously expressed *CLN3* we first designed a vector with the 5' UTR of *CLN3*, the designed *CLN3* sequences, a Kanamycin resistance gene, and then homology to the 3'

UTR of *CLN3*. We digested a previously published pR416 plasmid containing Cln3-GFP with AscI (New England Biolabs R0558S) and AatII (New England Biolabs R0117). This digest removed the Cln3 coding sequence and the GFP and resulted in a band of 7.5kb which was gel purified using the QIAquick gel extraction kit (Qiagen, 28704) according to the manufactures instructions. From the pJet Plasmids containing the *CLN3* sequences we amplified *CLN3* with primers that were universal to all sequences (fwd: tacctgcacgcggtcgagacgtctgcataccaaagatc, rev: tgcatttacttataatggttacttatcgtcatcgtccttgtagtcgcgtagtttcttgac). This PCR fragment was cloned into the backbone using the HiFi DNA assembly kit according to the manufacturer's instructions (New England Biolab E5520) and then transformed into 5-alpha competent cells (New England Biolabs, C2987). The resultant colonies were miniprepped using the GeneJet Kit (ThermoFisher K0502) according to the manufacturer's instructions and sequenced by long read and sanger to confirm the correct sequence (Azenta Life Science, Plainfield, NJ).
To transform *Ashbya* we generated a linear fragment containing the 5' UTR, the designed *CLN3* coding sequences, a kanamycin resistance gene and homology to the 3' UTR. This fragment was generated by PCR of the pRS416 vector (fwd: GGGCATGTTTGACACAATGG rev: CGGCTAGGAGACTGCATTTAT) resulting in a fragment of 3.9kb. Any remaining plasmid DNA was destroyed by the addition of DpnI (New England Biolabs R0176). Following this digest, the remaining DNA was purified and concentrated using the QIAquick PCR purification kit (Qiagen, 28104). This DNA was transformed into a previously published *Ashbya* strain that contains an endogenously tagged Whi3-tomato protein (*19*). Transformation was done using previously published protocols (*41*). Transformants were verified by PCR of the genomic DNA followed by Sanger sequencing (Genewiz, South Plainfield, NJ, USA).

Live cell imaging

Live imaging was based on previous published protocols (*42*). Briefly, cells were grown overnight in AFM containing G418-sulfate (gold bio 108321-42-2) at 30° in a baffled flask with gentle shaking (110rpm). Cells were sedimented by gentle centrifugation (200rpm for 4min) and then washed 2x with Low Fluorescence Media (LFM). After washing cells were plated on an agarose pad in a depression slide containing 1% (W/V) agarose in LFM, covered with a coverslip an imaged.

Imaging was performed with a Zeiss LSM 980 using an Apochromat 63x 1.40NA Oil immersion objective. The microscope was equipped with an incubation chamber allowing for control of temperature at 30°. Illumination was done using a 561nm diode laser and detection was with a high QE 32 channel spectral array GaAsP.

Scoring of Whi3 condensates in live cells

Individual hyphae were classified as belonging to one of three categories based on the appearance of their Whi3 signals: 1) big condensates – such hyphae contain only well-defined, large condensates with a dark dilute phase in the rest of the cytoplasm. 2) intermediate – such hyphae contain some large condensates but also some network-like Whi3 morphologies at the same time. 3) network – such hyphae contain only small, interconnected, network-like condensates throughout the entire cytoplasm. The total numbers of categorized hyphae for each *CLN3* mutant *Ashbya* strain are L1 (75), L2 (30), WT (17), H1 (60), and H2 (18).

Single Molecule RNA FISH

Single-molecule RNA FISH was conducted based on previously published protocols (*28, 43*). Cells were grown overnight in AFM at and then fixed in the flask with formaldehyde to a final concentration of 3.7% vol/vol for 1 h. The mycelia were gently pelted by centrifugation (400xcf) in a 15ml conical before being washed twice with ice-cold buffer B (1.2 M sorbitol and 0.1 M potassium phosphate, pH 7.5). The cells were resuspended in 1 ml spheroplasting buffer (10 ml buffer B and 2 mM vanadyl ribonucleoside complex) and transferred to a new 1.7ml RNase-free microcentrifuge tube. 1.5 mg Zymolase prepared with RNAse-free water was added to the cells and incubated at 37°C for 10 min to digest and permeabilize the cell walls. Digestion was monitored by DIC. following digestion cells were washed twice with buffer B to remove the zymolyase and then resuspended in 1 ml RNase-free 70% Ethanol overnight at 4°C.

Custom RNA FISH probes (Biosearch Technologies) were designed with homology to the coding sequence of each gene. Probes were labeled with TAMRA and were resuspended in 20 µl 1x TE buffer. The ethanol was removed from the cells and then they washed twice with wash buffer (2x SSC (0.3M Sodium Chloride, 30mM trisodium Citrate), 10% vol/vol deionized formamide), resuspended in 100 µl hybridization buffer (1 g dextran sulfate, 10 mg *E. coli* tRNA, 2 mM vanadyl ribonucleoside complex, 2 mg BSA, 2x SSC, and 10% vol/vol deionized formamide in 10mL final volume with RNAse-free water) with 25nM mRNA FISH probe. The cells were incubated with probe overnight in the dark at 37°C. Then cells were washed once with wash buffer. Then they were resuspended in wash buffer with Hoechst and incubated at 37°C for 30 min. Cells were then washed again with wash buffer, followed by two washes with RNAse free PBS. Cells were mounted with vectashield. covered with an RNase-free coverslip, sealed with nail polish, and imaged.

Imaging was performed with a Zeiss LSM 980 using a Apochromat 63x 1.40NA Oil immersion objective. Illumination was done using simultaneous illumination with 405nm and 561 nm diode lasers and detection was with a high QE 32 channel spectral array GaAsP with unmixing capability. Individual channels were resolved using the linear demixing function resulting in three channels for each image, a DAPI channel, a Fish Channel, and a channel that contained autofluorescence of the hyphae. Microscope control and analysis was performed on premium HP Z6 workstations running Zeiss ZEN 3.7.

Image segmentation was done in two steps. First each image was individually contrasted in ImageJ to maximize the SNR of each channel. Two images from each strain were chosen at random and cropped to include only a small subset of the hyphae. These were used as a training set for iLastic (*44*). In iLastic we trained the pixel classification model to recognize four categories of pixels: the area outside of the hyphae, the area inside the hyphae, nuclei and fish spots. Using these segments we measured the area of the hyphae and we measured both the number of FisH spots and calculated the integrated intensity of each spot. The total integrated intensity of most spots were normally distributed around a single value. We defined this peak as the intensity of one transcript, however a diffraction limited spot can contain multiple transcripts so any spot so any spot with double the median intensity was defined as two transcripts, and so forth.

Immunofluorescence

**Experimental assay:** Immunofluorescence was performed as described previously for *Ashbya* (*42*) based on a protocol for yeast (*45*). Briefly, cells were grown for 16hr at 30°C in a baffled flask with gentle shaking (110rpm). Cells were fixed with formaldehyde at a final concentration of 3.7% for 1hr while shaking. Cells were pelleted by centrifugation (400rcf) and washed 3x with PBS and then resuspended in Solution A (100mM K2PHO4, pH 7.5, 1.2M sorbitol) for digestion with zymolyse at 37°C to remove the cell wall. Digestion was monitored by phase contrast microscopy. Cells were blocked for 1hr in PBS with 1mg/ml BSA and 0.1% triton and then stained overnight with anti-alpha Tubulin antibody (1:100, YOL1/34 Bio-rad CAT# MCA78G; RRID: AB_93477). Cells were then washed, blocked again, and stained with secondary antibody (1:500, Goat-Anti-Rat conjugated to Cy3, Millipore-Sigma CAT # AP136C; RRID: AB_11211234) and Hoechst and then mounted with vectashield. Cells were imaged on a Nikon Ti-E stand equipped with a Yokogawa CSU-W1 spinning disk confocal unit, a Plan-Apochromat 100x/1.49 NA oil-immersion objective, and either a Prime 95b sCMOS camera (Photometrics) or a Zyla sCMOS camera (Andor). Nikon NIS-Elements software was v.4.60. The spinning disk was illuminated with separate 405 and 561 nm laser sources.

**Cell cycle state analysis:** All images with scored nuclei that were used in analysis are shown in **Fig S15**. Cell cycle states of each nucleus were aggregated for each *Ashbya* strain and plotted as stacked bar plots in **Fig 3G**.

**Nuclear synchrony analysis:** The same images shown in **Fig S15** were used for nuclear synchrony analysis. All pairs of consecutive, scored nuclei within the same hypha were used in analysis. The synchrony index was calculated as described in (*28*). Briefly, the observed synchrony is divided by the chance synchrony, which gives values greater than 1 if nuclear divisions are more synchronous than chance, and less than 1 if they are more asynchronous than expected by chance. Calculation of the index is implemented in a google sheet: https://docs.google.com/spreadsheets/d/146DXA55su_luV6xismBkRsT5sGqmgGKe6S4WBUWy0_g/edit#gid=0. Results are shown in **Table S3**.

Hyphal growth rate assay

FITC-ConA staining was performed as previously described (*46*). *Ashbya* were grown from dirty spores in 10 mL AFM for 14 hours at 30ºC. Cell pellets were collected by centrifuging cultures for 5 minutes at 300 rpm in 15 mL tubes. Cells were then resuspended and washed twice in 10 mL wash buffer containing 50 mM Tris pH 7.5,150 mM NaCl. 50 µL of 1 mg/mL FITC-ConA stock was added to cell pellet resuspended in 2 mL of the wash buffer and shaken at room temperature for 10 minutes. Cells were washed 2X with 10 mL AFM, resuspend cells in 10 mL AFM and appropriate selection. They were allowed to grow in a baffled flask at 300 rpm at 30°C for 1 h before fixation with 1 mL 37% formaldehyde for 1 h, shaking. Cells were washed with 1X PBS, resuspended in 500 µL 1X PBS and stained with Hoechst before mounting in 10 µL Vectashield mounting medium (Vector Laboratories H-1000-10). FITC-ConA signal was imaged using a spinning disc confocal, and growth rates were measured by measuring the length of the tip region unlabeled with FITC in Fiji.

### Supplementary Text

Relationship between ensemble diversity and the variance of the nucleotide structural state

The following defines a mathematical relationship between the ensemble diversity and the variance of the ensemble probability that a nucleotide is single-stranded for a given sequence. The distribution of these probabilities is over all of the nucleotides in the sequence which generates the structural ensemble. We define $p_{ij}$ as the ensemble-level base pairing probability between nucleotides $i$ and $j$, $p_i^* = 1 - \sum_j p_{ij}$ as the probability of nucleotide $i$ being single-stranded, and $\mu^* = \frac{1}{N}\sum_i p_i^*$ as the average fractional single-stranded content for a sequence of length $N$. We first write the ensemble diversity (ED) in a more convenient form:

$$\mathrm{ED} = \sum_i \sum_j p_{ij}(1 - p_{ij}) = \sum_i \left( \sum_j p_{ij} - \sum_j p_{ij}^2 \right) = \sum_i \left( 1 - p_i^* - \sum_j p_{ij}^2 \right) =$$

$$N - N\mu^* - \sum_i \sum_j p_{ij}^2 \quad \rightarrow \quad \frac{\mathrm{ED}}{N} = -\frac{1}{N} \sum_i \sum_j p_{ij}^2 - \mu^* + 1$$

We want to understand how the variance of the $p_i^*$ is related to the ED. We begin with the definition of the variance:

$$\mathrm{Var}(p_i^*) = \frac{1}{N}\left( \sum_i (p_i^* - \mu^*)^2 \right) = \frac{1}{N}\left( \sum_i ((p_i^*)^2 - 2\mu^* p_i^* + (\mu^*)^2) \right) =$$

$$\frac{1}{N}\left( \sum_i (p_i^*)^2 - 2N(\mu^*)^2 + N(\mu^*)^2 \right) = \frac{1}{N}\sum_i (p_i^*)^2 - (\mu^*)^2 = \frac{1}{N}\sum_i \left( 1 - \sum_j p_{ij} \right)^2 - (\mu^*)^2$$

Expanding the term inside the sum over $i$:

$$\left( 1 - \sum_j p_{ij} \right)^2 = (1 - p_{i1} - p_{i2} - \cdots - p_{iN})(1 - p_{i1} - p_{i2} - \cdots - p_{iN}) =$$

$$1 - 2\sum_j p_{ij} + p_{i1}^2 + p_{i2}^2 + \cdots + p_{iN}^2 + 2\sum_k \sum_{j \neq k} p_{ik} p_{ij} =$$

$$\sum_j p_{ij}^2 + 2\sum_k \sum_{j \neq k} p_{ik} p_{ij} + 2\left( 1 - \sum_j p_{ij} \right) - 1 = \sum_j p_{ij}^2 + 2\sum_k \sum_{j \neq k} p_{ik} p_{ij} + 2p_i^* - 1$$

Plugging in, we have the following expression for the variance of the $p_i^*$:

$$\mathrm{Var}(p_i^*) = \frac{1}{N}\sum_i \left( \sum_j p_{ij}^2 + 2\sum_k \sum_{j \neq k} p_{ik} p_{ij} + 2p_i^* - 1 \right) - (\mu^*)^2 =$$

$$\frac{1}{N}\sum_i \sum_j p_{ij}^2 + \frac{2}{N}\sum_i \sum_k \sum_{j \neq k} p_{ik} p_{ij} - (\mu^*)^2 + 2\mu^* - 1$$

We can now rewrite the variance of the $p_i^*$ as a function of the ED:

$$\mathrm{Var}(p_i^*) = -\frac{1}{N}\mathrm{ED} + \frac{2}{N}\sum_i \sum_k \sum_{j \neq k} p_{ik} p_{ij} + \mu^*(1 - \mu^*)$$

We therefore expect $\mathrm{Var}(p_i^*)$ to be a linear function of the ED with a negative slope. Indeed, we see that this expectation is met both for the structural ensemble prediction by RNAfold and for the single molecule structure probing data (**Fig S8**).

Physical Information

The concept of physical information can be explained by the connections between ensemble diversity, Shannon entropy, and conformational entropy. The ensemble diversity can be interpreted analogously to the Shannon entropy, a central quantity in information theory, and is most easily understood when comparing their mathematical definitions. The ensemble diversity of a sequence $S$ with $N$ nucleotides is defined as $ED(S) = \sum_i p_i(1 - p_i)$, where $p_i$ is the probability of observing nucleotide $i$ in a structured state, given a structural ensemble generated by sequence $S$. The Shannon entropy of a random variable $X$ is defined as $H(X) = -\sum_{x \in X} p(x) \log p(x)$, where $p(x)$ is the probability of observing outcome $x$. Both quantities increase monotonically as the set of $p_i$ or $p(x)$ approaches a uniform distribution, and they both approach 0 as the probability of observing a single outcome goes to 1 (**Fig S17A**). We also propose that the ensemble diversity of sequence $S$ should scale with the conformational entropy as long as the conformational entropy is not too high (**Fig S17B**). This relationship can be seen by considering the behavior of an RNA sequence with increasing amounts of average intra-molecular base pairing. In the limit of a sequence in which all nucleotides are base-paired, the ensemble diversity is 0 since only 1 secondary structure is possible, and the conformational entropy is also 0 since no spatial sampling is possible. As the average content of unpaired nucleotides in the structural ensemble increases, more secondary structures become possible, and both the ED and the conformational entropy increase. However, once the average single-stranded content increases above a critical value, the ED will begin to decrease while the conformational entropy continues to increase. In the limit of a sequence in which no nucleotides are base-paired, the ED is again 0 since there is only 1 secondary structure, but the conformational entropy is maximized since the chain will have unrestricted access to all spatial arrangements. We argue that our sequences are below this critical value because the average unpaired nt. does not vary significantly with ED (**Fig S18**). Overall, we can see that the ED represents a physical information, in which a high ED corresponds both to high conformational and Shannon entropies.

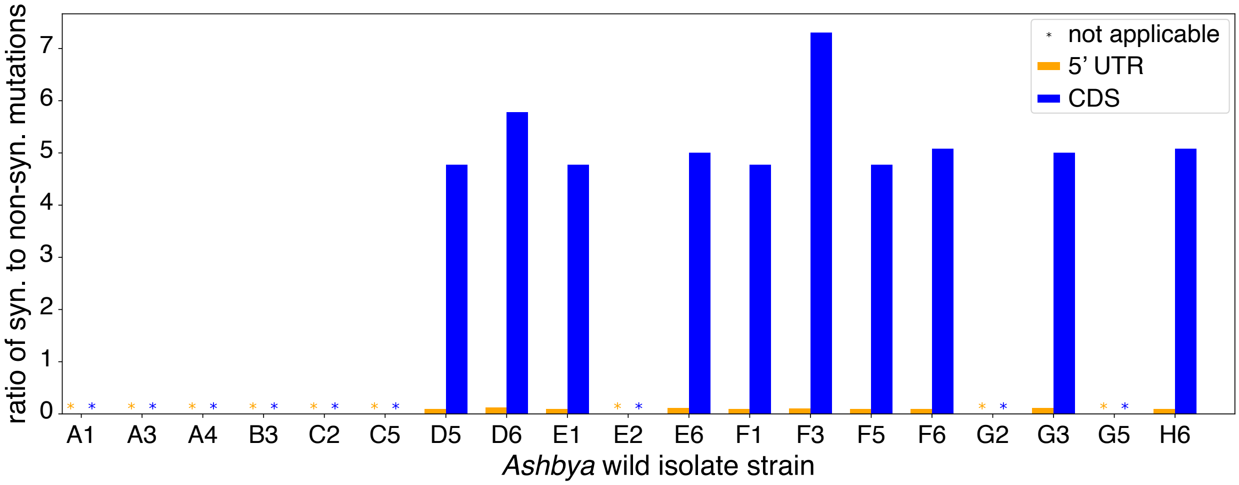


**Fig. S1: Ratios of synonymous to non-synonymous mutations in the 5′ UTR and CDS of *CLN3* from *Ashbya* wild isolates.** Xtick labels indicate the name of the *Ashbya* wild isolate strain. The lab reference strain is A1.

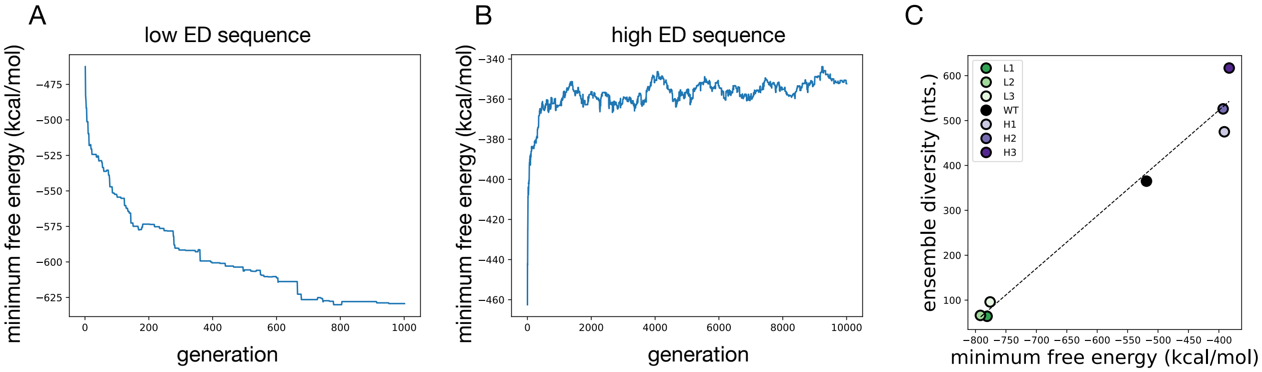


**Fig. S2: *in silico* evolution of *CLN3* structure mutants, and MFE-ED correspondence. (A)** The minimum free energy (MFE) of the chosen sequence in each generation is shown for an example evolution of a low ED sequence. **(B)** Equivalent to **(A)**, but for evolution of a high ED sequence. **(C)** The MFE and ED, calculated at a temperature of 25C and salt concentration of 150mM, are shown for each *CLN3* structure mutant in this study.

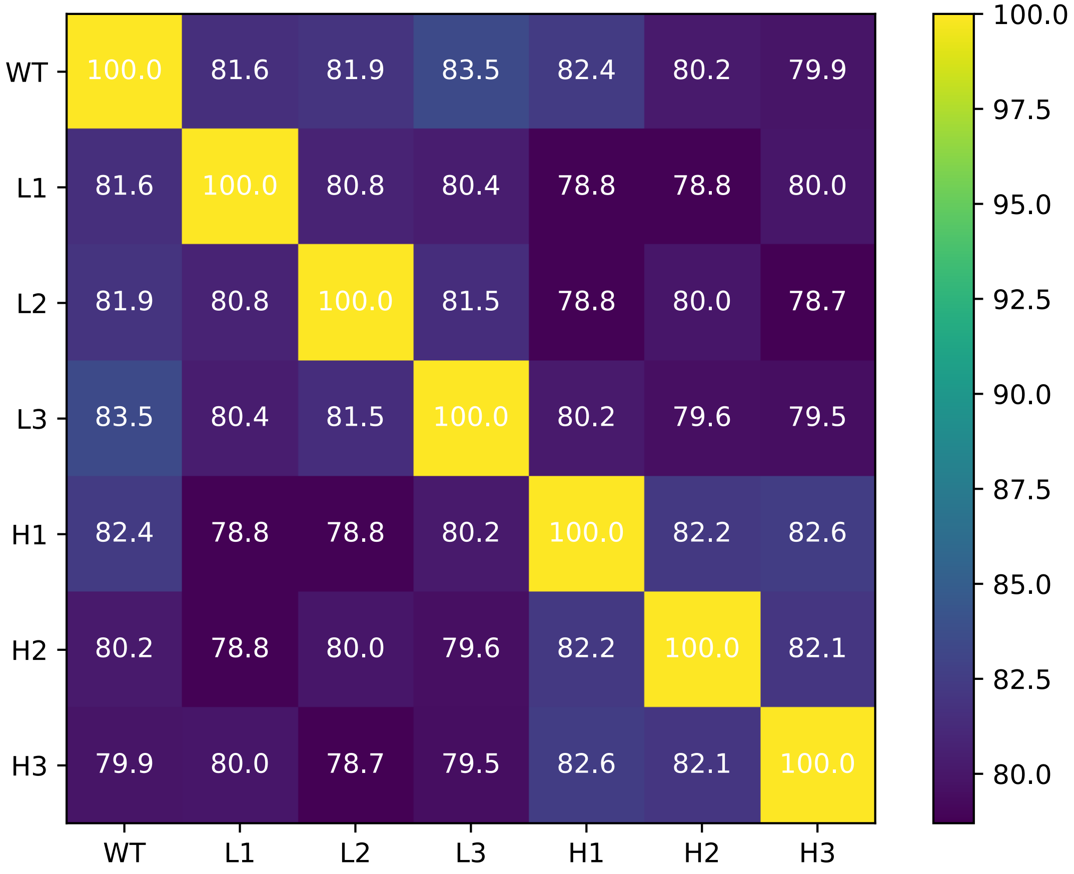


**Fig. S3: Sequence identity among *CLN3* structural mutants.** White numbers indicate the percentage sequence similarity among all *CLN3* sequences studied. Bin color indicates percentage similarity.

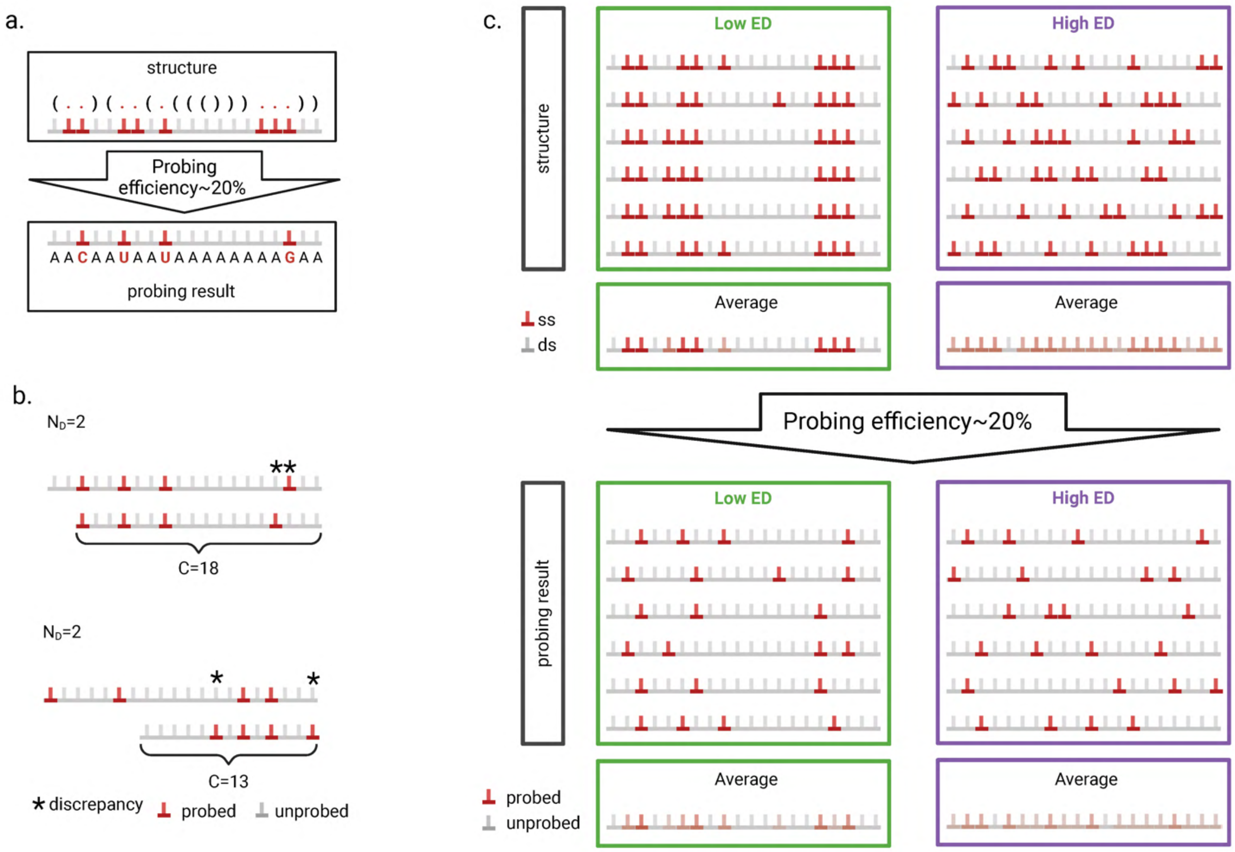


**Fig. S4: Structure probing and calculation of pairwise distances among reads. (A)** Schematic that demonstrates how single stranded adenosine are partially probed and reflected as mismatches after alignment. **(B)** Examples of how discrepancy and coverages are extracted to calculate pairwise distance. **(C)** Schematic demonstrating the low ED structures are more similar to each other than high ED, which results in more concentrated single-stranded nucleotides. This difference is dampened by imperfect probing efficiency.

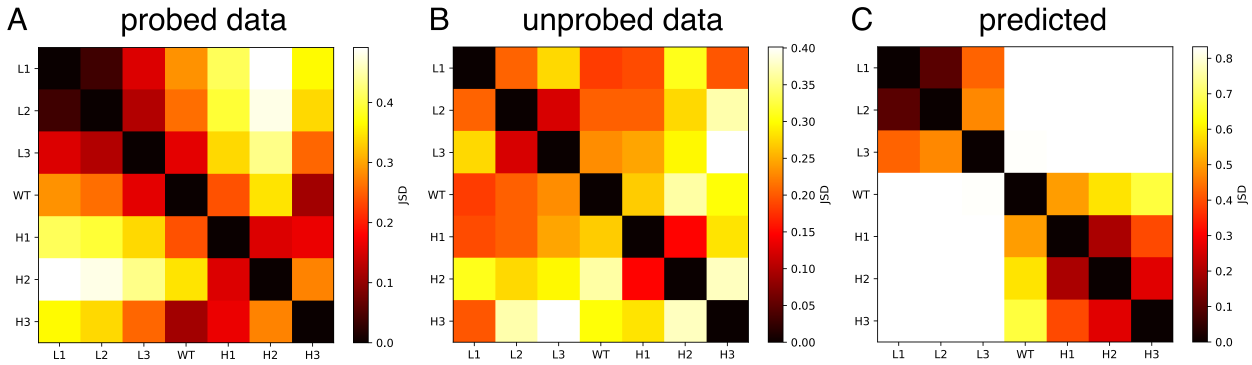


**Fig. S5: Jensen-Shannon distance between all normalized pairwise distance distributions. (A)** The Jensen-Shannon distance (JSD) among all of the normalized pairwise distance distributions shown in **Fig 1D** are shown as a heatmap with a JSD of 0 corresponding to black and the maximum JSD in the set corresponding to white. **(B)** The JSD among all of the normalized pairwise distance distributions from the unprobed experimental data are shown. The average JSD among the low ED pairwise distance distributions is 0.24, among the high ED distributions is 0.29, and between the low and high ED distributions is 0.31. **(C)** The JSD among all of the normalized pairwise distance distributions from the Boltzmann sampled predictions for each sequence are shown. The average JSD among the low ED pairwise distance distributions is 0.34, among the high ED distributions is 0.33, and between the low and high ED distributions is 0.83.

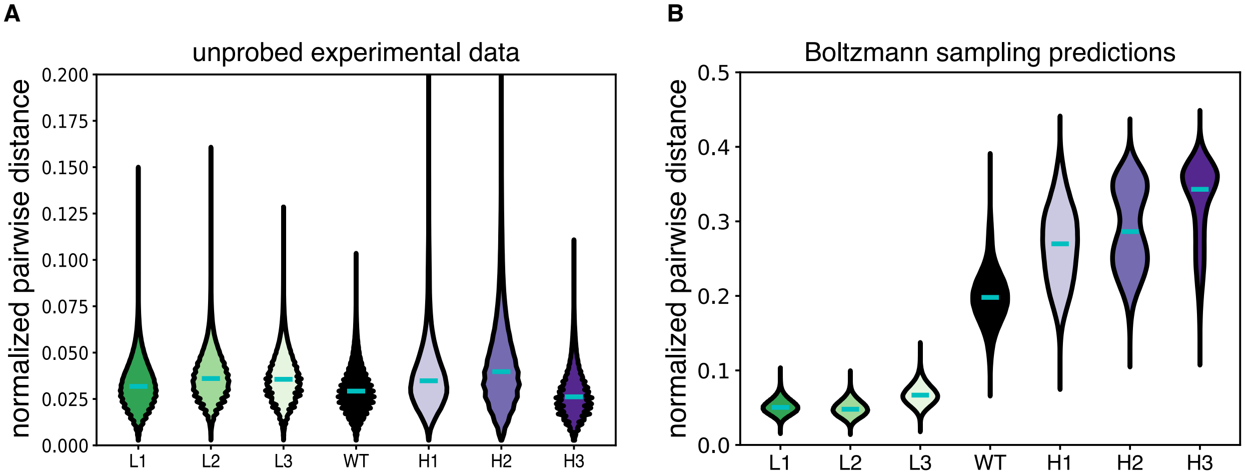


**Fig. S6: Normalized pairwise distance distributions for unprobed experimental data and RNAfold predictions. (A)** Normalized pairwise distances are computed for each sequence using Nanopore sequencing without addition of the probe molecule DEPC. Differences among sequences are no longer apparent as expected. **(B)** For each *CLN3* structural mutant, 1000 structures are sampled from the structural ensemble using Boltzmann sampling in the ViennaRNA package. The normalized pairwise distances are computed analogously as for the experimental structure probing data which is plotted in **Fig. 1D**. Cyan bars represent medians.

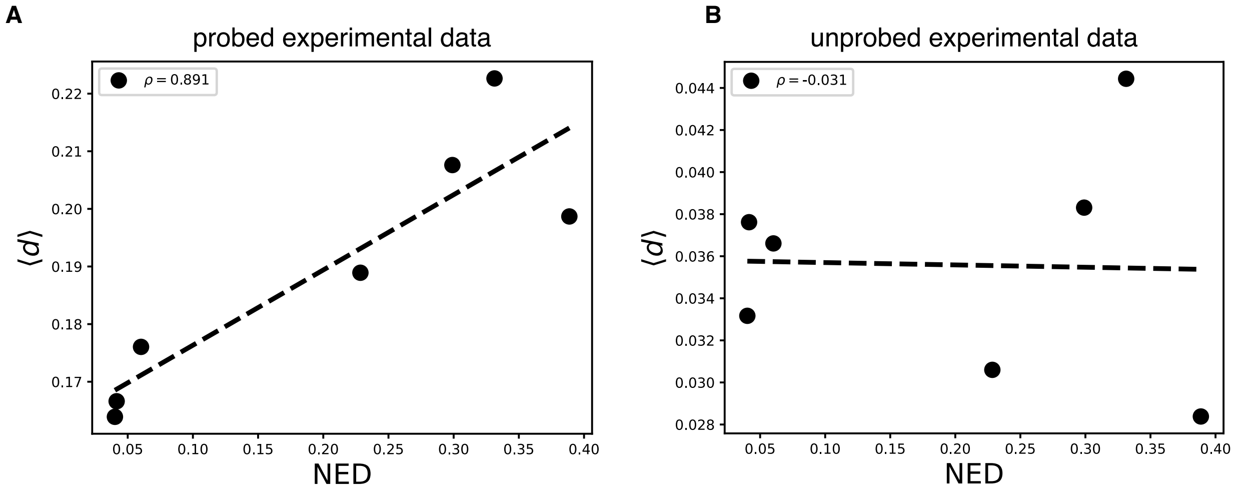


**Fig. S7: Pearson's correlation between predicted length-normalized ED and experimentally measured average pairwise distance. (A)** Data corresponds to DEPC probed experimental data shown in **Fig. 1D**. $\rho = 0.891$ is the Pearson's correlation between $\langle d \rangle$ and NED. **(B)** Data corresponds to an experimental control without DEPC probing. $\rho = -0.031$ is the Pearson's correlation between $\langle d \rangle$ and NED.

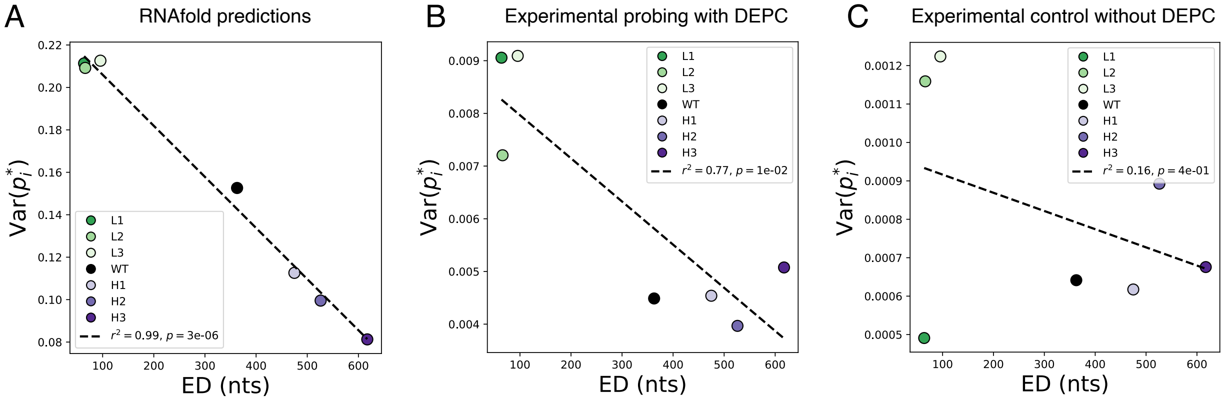


**Fig. S8: Scaling of the variance of the per-nucleotide probability of being single-stranded, $p_i^*$, with ensemble diversity. (A)** For each sequence, 10,000 structures were generated using Boltzmann sampling in RNAfold at 25ºC and 150 mM NaCl, and $p_i^*$ for each adenosine nucleotide was calculated. The variance of the $p_i^*$ is plotted against the ensemble diversity (ED) of each sequence calculated at 25ºC and 150 mM NaCl. The dotted line is a linear regression with a coefficient of determination of 0.99 and a p-value of $3 \times 10^{-6}$. **(B)** The $p_i^*$ for each adenosine for which there were at least 100 reads from the single molecule structure probing experiments corresponding to **Fig. 1C** was calculated for each sequence, and the variance of the $p_i^*$ is plotted against the predicted ED calculated at 25ºC and 150 mM NaCl. The dotted line is a linear regression with a coefficient of determination of 0.77 and a p-value of 0.01. **(C)** The single molecule structure probing experiment was repeated without the chemical probe DEPC as a negative control. The $p_i^*$ for each adenosine for which there were at least 100 reads for each sequence was calculated, and the variance of the $p_i^*$ is plotted against the predicted ED calculated at 25ºC and 150 mM NaCl. The dotted line is a linear regression with a coefficient of determination of 0.16 and a p-value of 0.4.

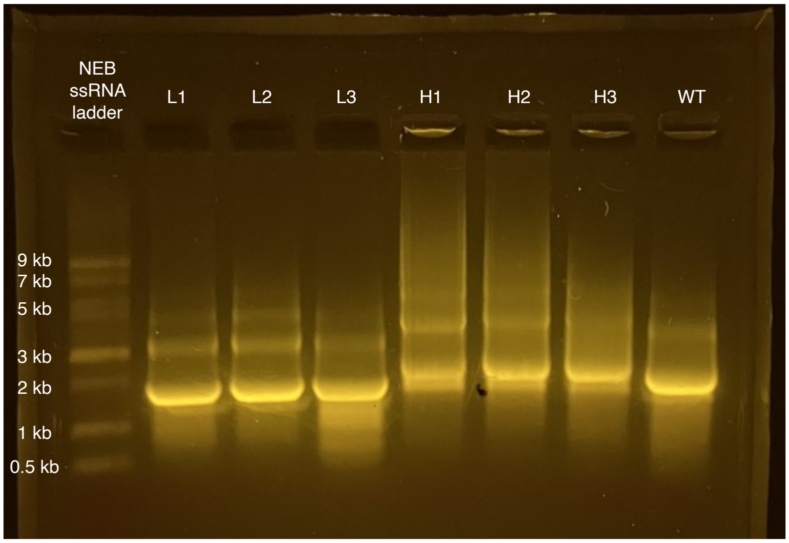


**Fig. S9: Native RNA gel.** 2 μM of each RNA in 150 mM KCl 50 mM HEPES pH 7.4 (phase separation buffer) was loaded into a 1% agarose gel in TBE (89 mM Tris-HCl pH 7.8, 89 mM borate, 2 mM EDTA) with a 2:50000 volume fraction of SYBR Safe DNA gel stain (Thermo Fisher).

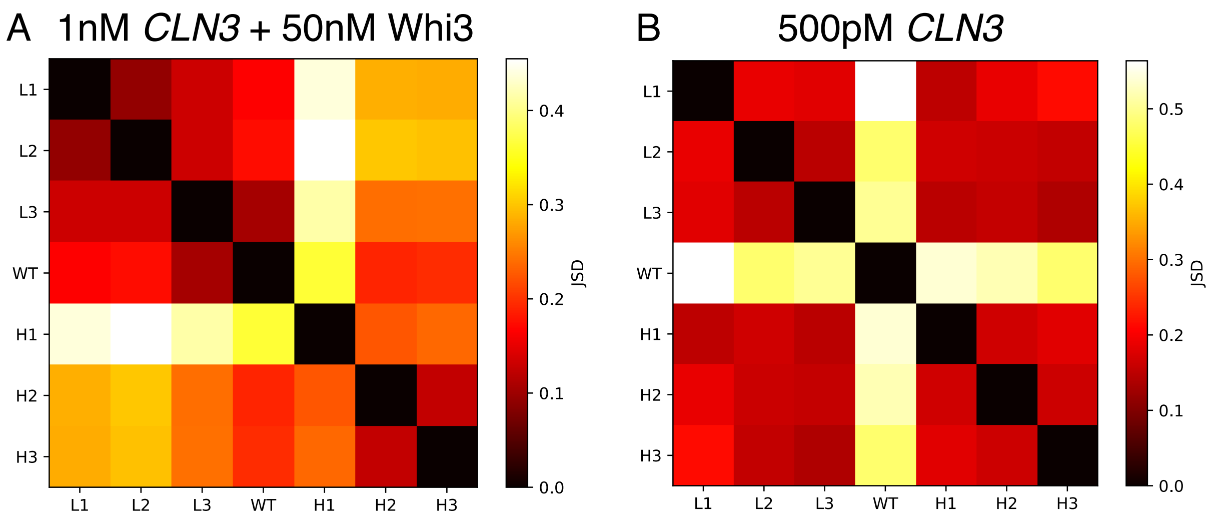


**Fig. S10: Jensen-Shannon distance (JSD) for subsaturated TIRF experiments. (A)** The JSDs among all of the *CLN3* intensity distributions shown in **Fig 2B** and corresponding to experiments with 1nM *CLN3* mixed with 50nM Whi3 are indicated with the heatmap. **(B)** The JSDs among all of the *CLN3* intensity distributions shown in **Fig S11** and corresponding to experiments with only 500pM *CLN3* are indicated with the heatmap. The average JSD among the low ED *CLN3* intensity distributions is 0.17, among the high ED distributions is 0.17, and between the low and high ED distributions is 0.16.

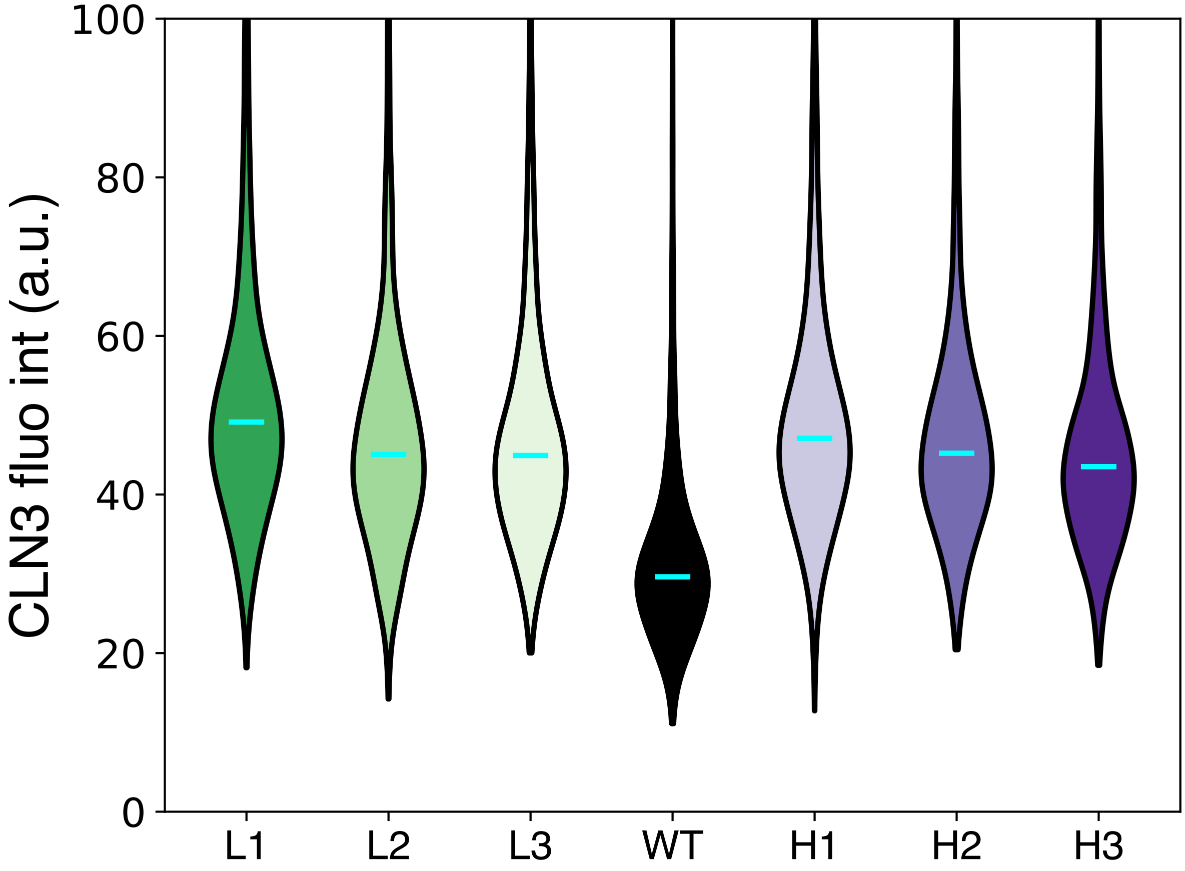


**Fig. S11: TIRF microscopy for each RNA without added protein.** 500 pM of each cy5-labeled RNA was incubated in 150 mM KCl 50 mM HEPES pH 7.4 (phase separation buffer) for 1 hour at 25ºC, pipetted onto plasma-cleaned glass, and imaged with TIRF microscopy. The average intensity of each puncta was extracted and the distributions for each sequence are shown as violin plots. Cyan bars indicate medians.

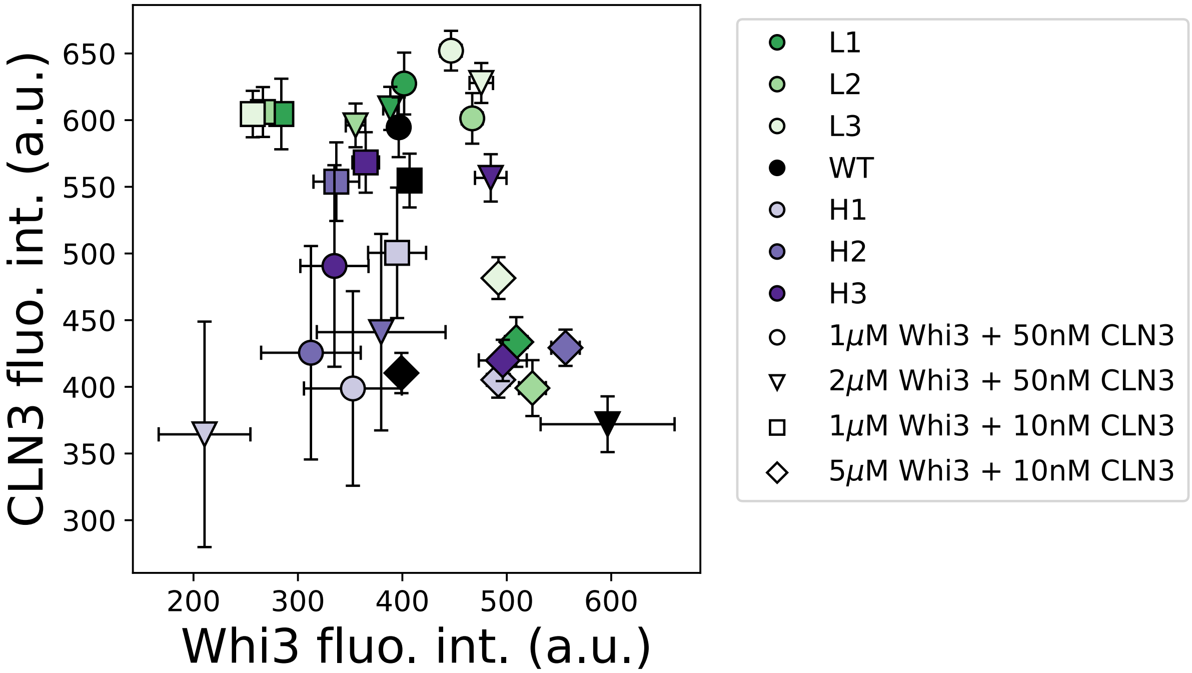


**Fig. S12: Dense phase concentrations of all *CLN3* structure mutants at four different (Whi3, *CLN3*) initial concentrations.** Mean dense phase fluorescence intensities of all *CLN3* + Whi3 systems are shown. All fluorescence intensities have been divided by 100. Colors indicate the *CLN3* structural mutant and marker shapes indicate the initial concentrations of Whi3 and *CLN3* used in the experiment. Error bars indicate standard deviations.

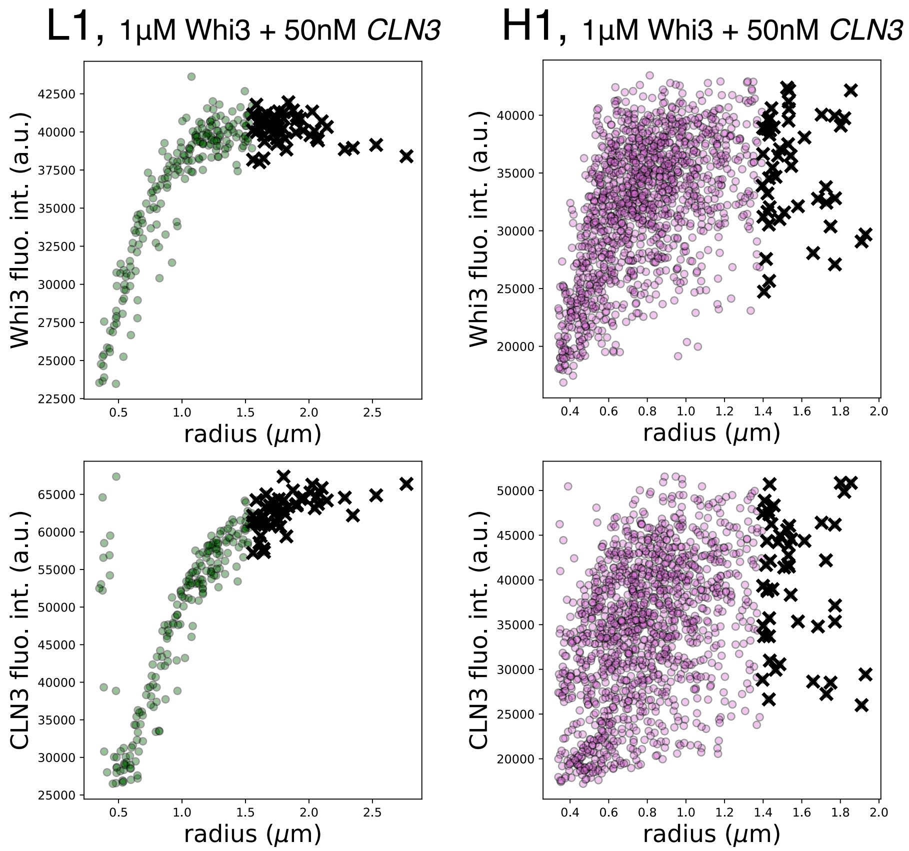


**Fig. S13: Condensate radius versus fluorescence intensity and selected observations for phase diagrams.** In each panel, every colored, circular marker corresponds to the radius of a single condensate and the average fluorescence intensity of the 27 pixels centered at the centroid of the same condensate. The black x's mark the 50 condensates with the largest radii in each set whose values are used in further analysis. Each panel corresponds to all condensates in 5 separate images corresponding to the same condition. The representative conditions shown are the L1 and H1 systems at the initial concentrations of 1 µM Whi3 + 50 nM *CLN3*, the results of which are shown in the phase diagram in **Fig. 3B** in the main text.

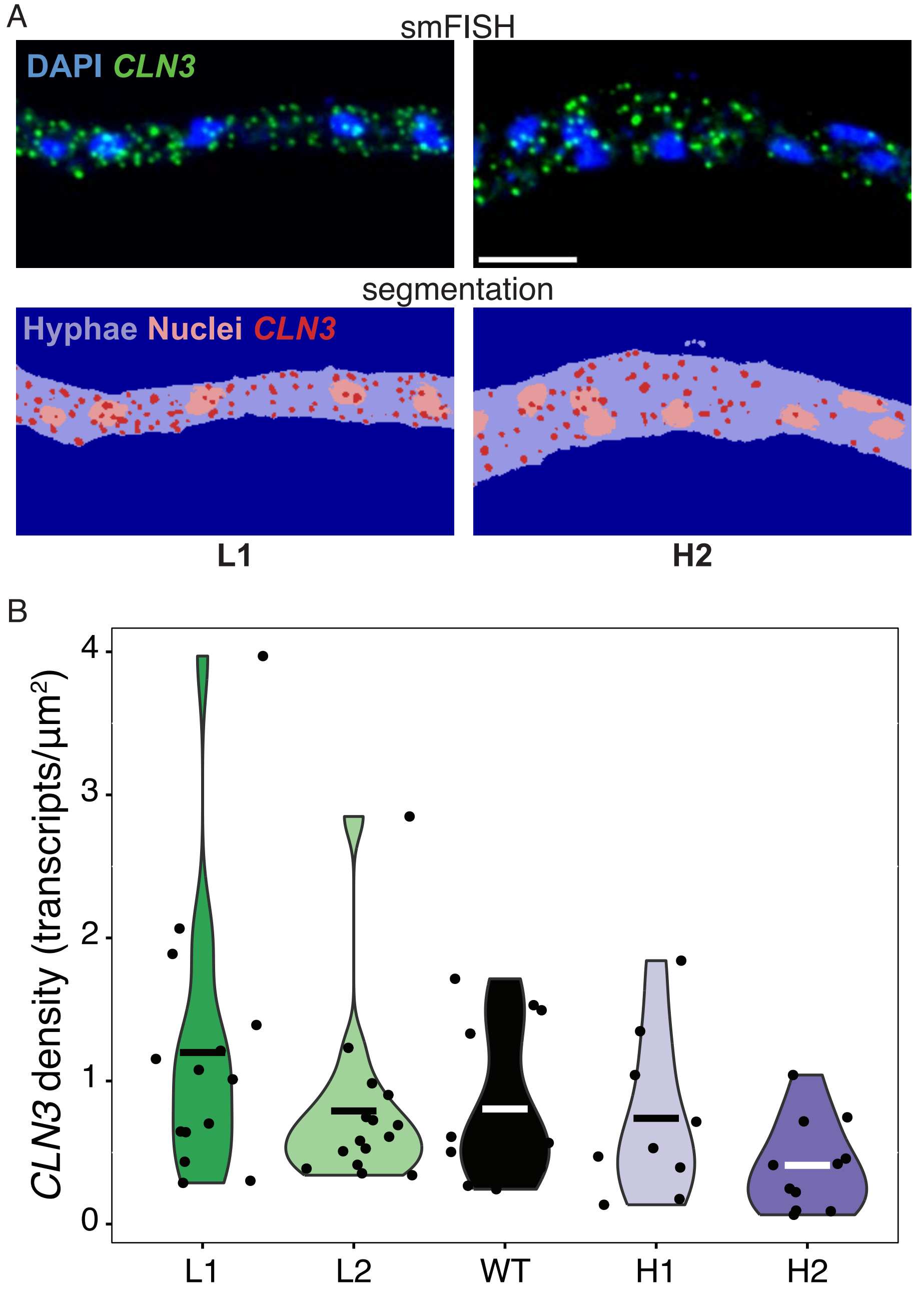


**Fig S14: *In vivo* CLN3 concentration measured by single molecule RNA Fluorescent *in situ* Hybridization (smFISH). (A)** Representative images of smFish from a L1 and H2 hyphae, showing the original image (top) and the segmented images (bottom). **(B)** Quantification of the number of *CLN3* molecules per µm$^2$ of hyphae. Dots represent individual hyphae, bars represent means of distributions.

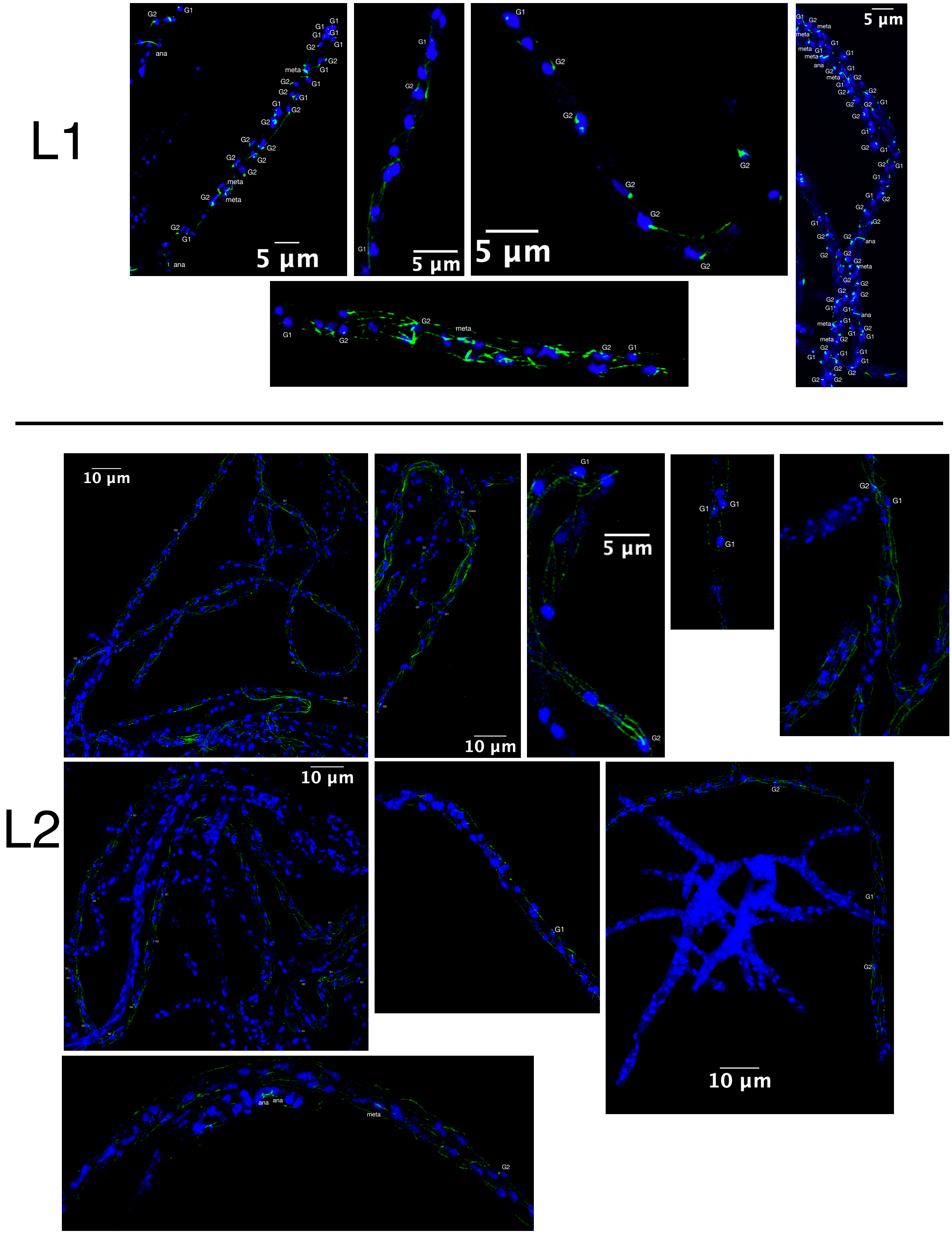


**Fig. S15: Immunofluorescence staining of *Ashbya* to determine cell cycle states.** Figure continued on next page.

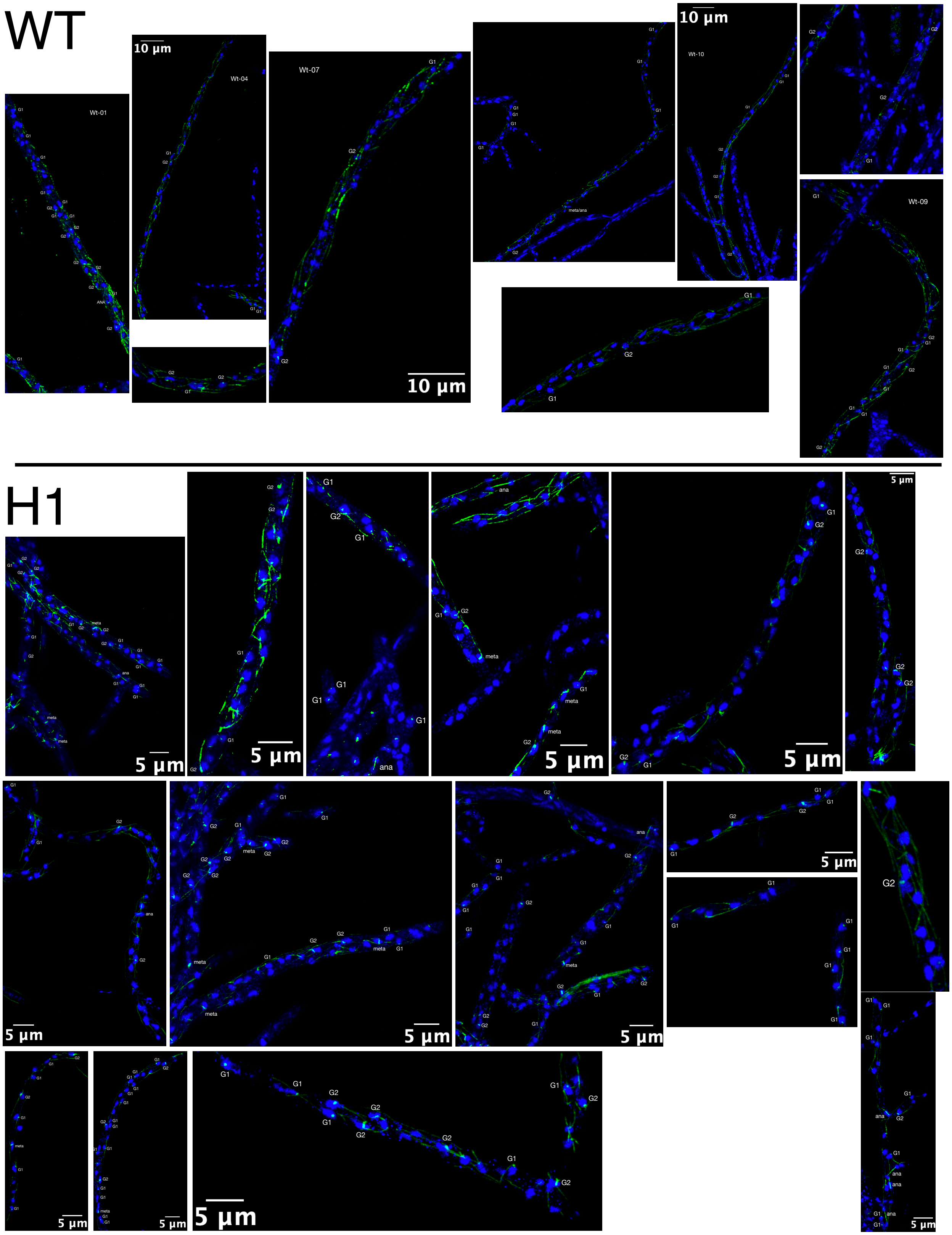


**Fig. S15: Immunofluorescence staining of *Ashbya* to determine cell cycle states.** Figure continued on next page.

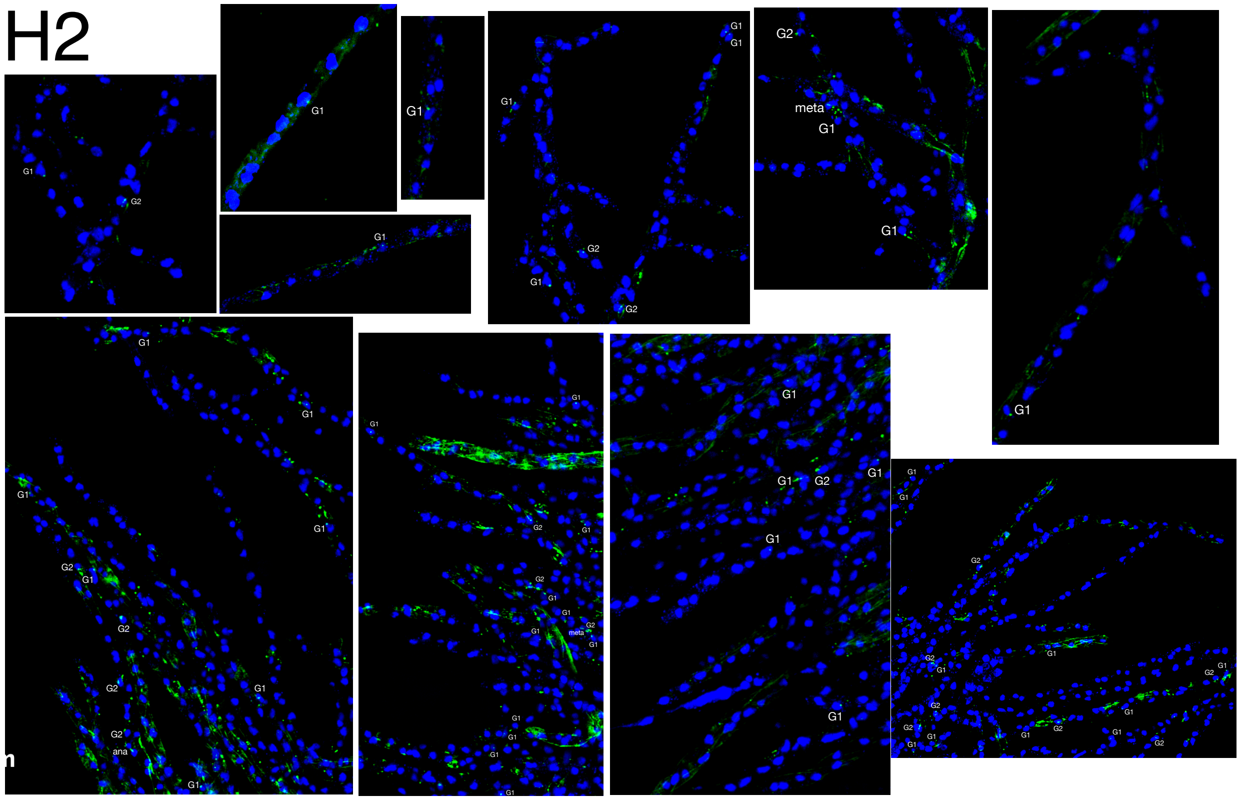


**Fig. S15: Immunofluorescence staining of *Ashbya* to determine cell cycle states.** All images used in analysis of cell cycle state (**Fig. 3G**) and nuclear synchrony (**Table S3**) are shown. Nuclei are shown in blue and spindle pole bodies are shown in green.

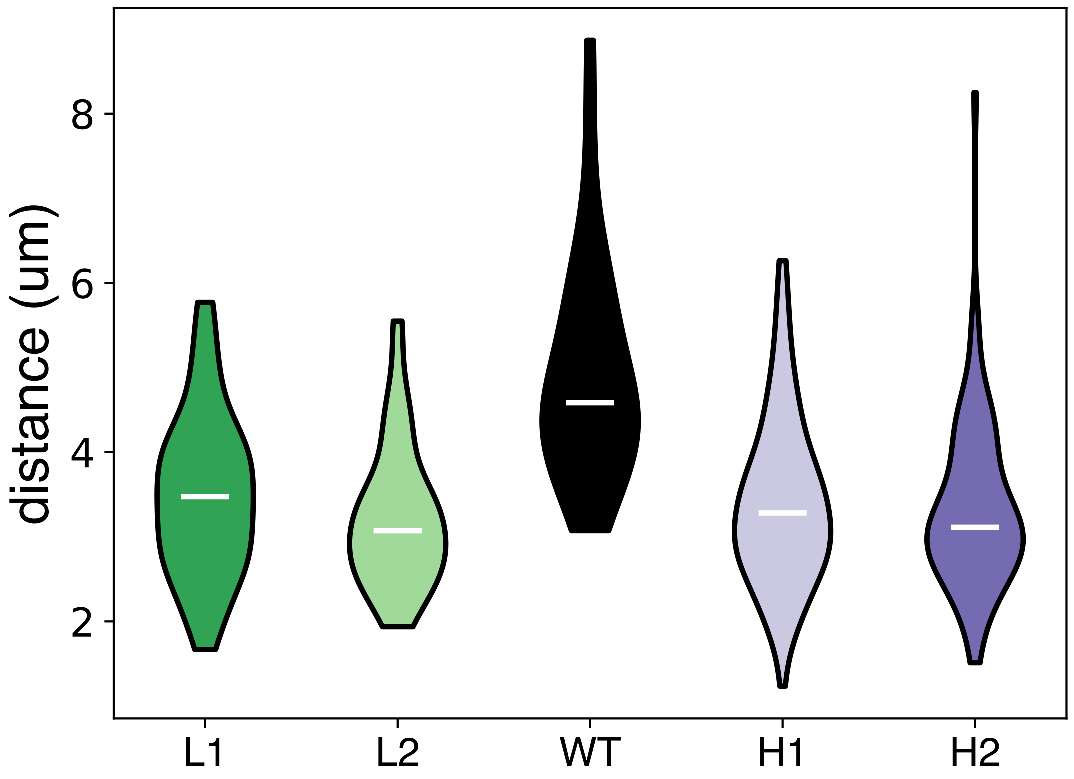


**Fig. S16: Internuclear distances in the *CLN3* structure mutant *Ashbya* strains.** Distributions of internuclear distances are shown for each strain. White bars indicate medians.

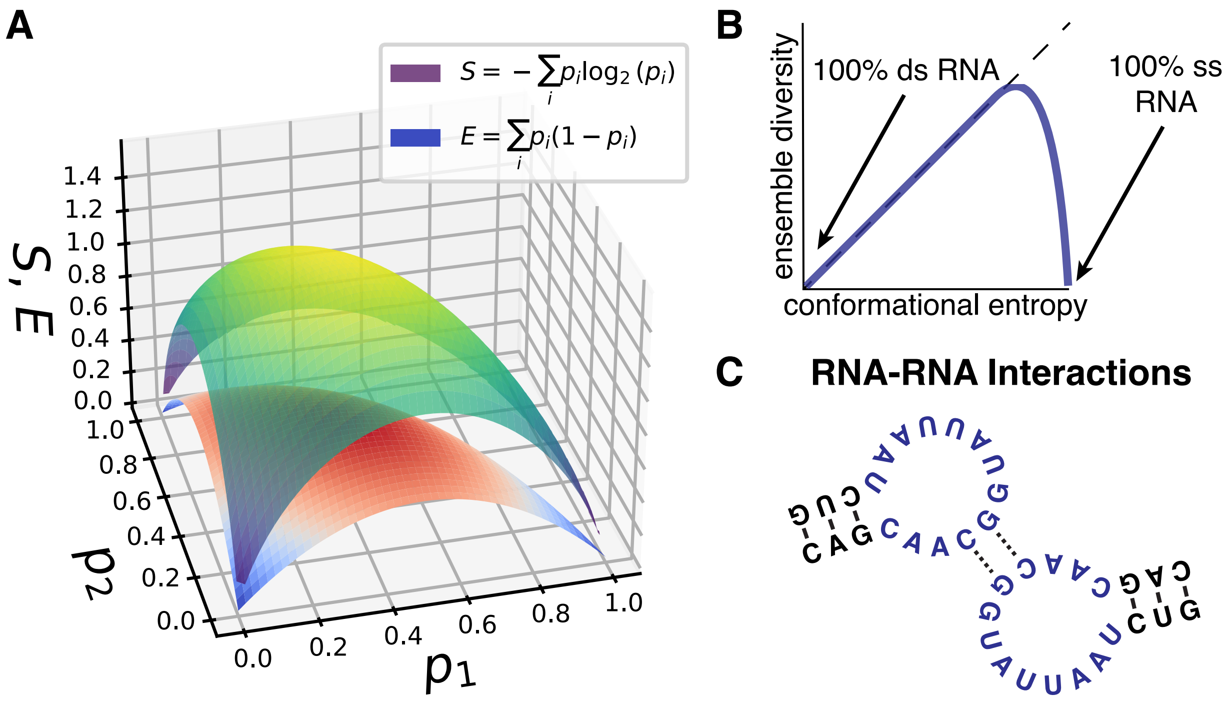


**Fig. S17: Connections between ensemble diversity, Shannon entropy, and conformational entropy. (A)** The Shannon entropy, S (purple-green-yellow gradient), and the ensemble diversity, E (blue-red gradient), are shown as a function of $p_1$ and $p_2$ in a system with 3 degrees of freedom. **(B)** Ensemble diversity is related to conformational entropy as shown in the schematic graph. ED as a function of conformational entropy is plotted as a thick purple line, and the identity relation is plotted as a dotted line. "ds" indicates double-stranded, and "ss" indicates single-stranded. **(C)** RNA sequences encode RNA-RNA interactions which depend on both sequence complementarity and RNA structural ensembles, since only nucleotides which are not participating in intra-molecular bonds (purple) are available for inter-molecular interactions.

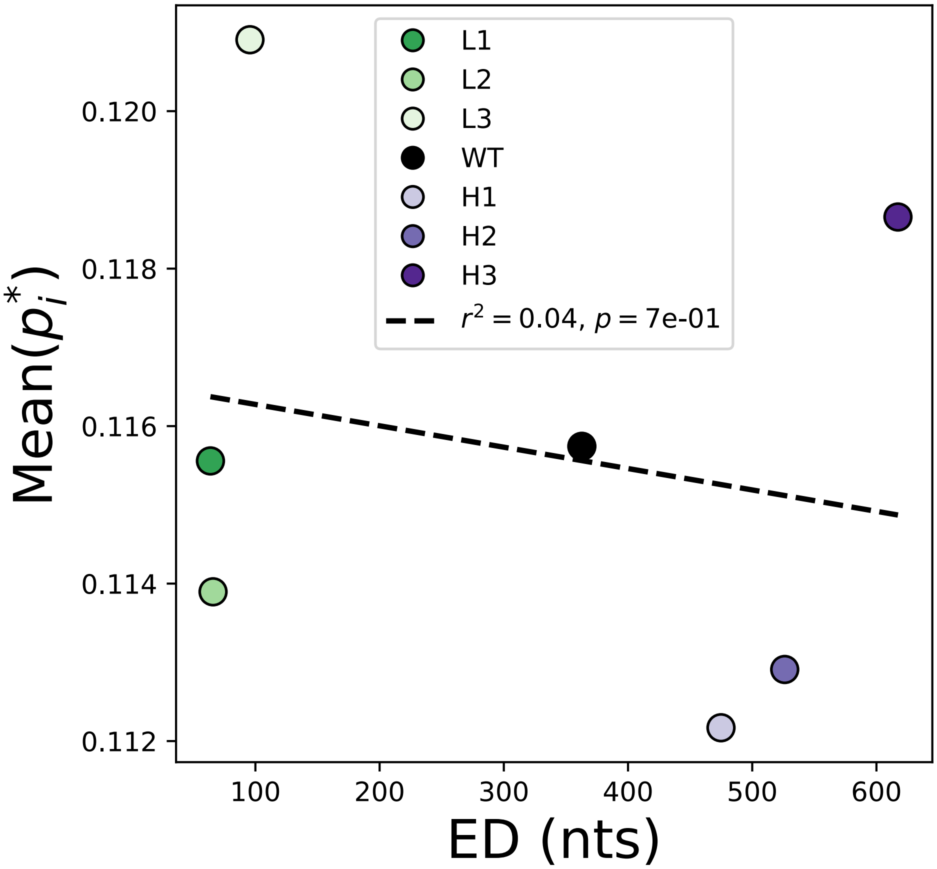


**Fig. S18: Average single-stranded content determined by experimental structure probing versus ensemble diversity.** The colored markers show the average single-stranded content of each *CLN3* structure mutant as determined in the single-molecule structure probing experiments. The dotted line is the best linear fit, $r^2$ indicates the goodness of fit, and p is the associated p-value.

| *CLN3* structure mutant | 3' extension sequence |
|---|---|
| L1 | GUUCCUUCCG |
| L2 | CUCUUCUCUU |
| L3 | UUGGUUCUCG |
| WT | GUUCCUGUGC |
| H1 | GUGUCGGUUG |
| H2 | GUCUGUGGUG |
| H3 | GCCUCUGUCU |

**Table S1: 3' extension sequences used in Nanopore sequencing.** The indicated sequences were added to the 3' end of each RNA for use in the Direct-RNA Nanopore chemical probing experiments.

| oligo | sequence |
|---|---|
| A | /5PHOS/GGCTTCTTCTTGCTCTTAGGTAGTAGGTTC |
| L1-B | GAGGCGAGCGGTCAATTTTCCTAAGAGCAAGAAGAAGCC(CGGAAGGAAC) |
| L2-B | GAGGCGAGCGGTCAATTTTCCTAAGAGCAAGAAGAAGCC(AAGAGAAGAG) |
| L3-B | GAGGCGAGCGGTCAATTTTCCTAAGAGCAAGAAGAAGCC(CGAGAACCAA) |
| WT-B | GAGGCGAGCGGTCAATTTTCCTAAGAGCAAGAAGAAGCC(GCACAGGAAC) |
| H1-B | GAGGCGAGCGGTCAATTTTCCTAAGAGCAAGAAGAAGCC(CAACCGACAC) |
| H2-B | GAGGCGAGCGGTCAATTTTCCTAAGAGCAAGAAGAAGCC(CACCACAGAC) |
| H3-B | GAGGCGAGCGGTCAATTTTCCTAAGAGCAAGAAGAAGCC(AGACAGAGGC) |

**Table S2: A and B oligo sequences for Nanopore sequencing library preparation.** Oligo A was used for every sequence while the oligo B sequences were RNA-specific.

| *Ashbya* strain | Synchrony Index | p-value | Nuclei-nuclei count |
|---|---|---|---|
| L1 | 1 | 0.494 | 70 |
| L2 | Insufficient data | Insufficient data | 4 |
| WT | 1.1 | 0.373 | 18 |
| H1 | 1.01 | 0.487 | 123 |
| H2 | 0.5 | 0.135 | 10 |

**Table S3: Nuclear synchrony analysis.** The synchrony index and associated p-value are shown and calculated as described in Methods section Immunofluorescence. The nuclei-nuclei count column indicates the number of neighboring nuclei used in the calculation of the synchrony index for each *Ashbya* strain.

**Movie S1.**
Several hyphae from the *Ashbya* strain with the L1 *CLN3* structural mutant integrated into the genome were imaged every 10s for 10 minutes with a 561nm laser to illuminate Whi3-tdTomato. The black excluded circular regions within hyphae are nuclei.

**Movie S2.**
One hypha from the *Ashbya* strain with the L2 *CLN3* structural mutant integrated into the genome was imaged every 10s for 10 minutes with a 561nm laser to illuminate Whi3-tdTomato. The black excluded circular regions within hyphae are nuclei.

**Movie S3.**
Several hyphae from the *Ashbya* strain with the H1 *CLN3* structural mutant integrated into the genome was imaged every 10s for 2 minutes with a 561nm laser to illuminate Whi3-tdTomato. The black excluded circular regions within hyphae are nuclei.

**Movie S4.**
Two hyphae from the *Ashbya* strain with the H2 *CLN3* structural mutant integrated into the genome was imaged every 10s for 5 minutes with a 561nm laser to illuminate Whi3-tdTomato. The black excluded circular regions within hyphae are nuclei.
